# Neurofibromin 1 controls metabolic balance and Notch-dependent quiescence of juvenile myogenic progenitors

**DOI:** 10.1101/2021.09.28.462168

**Authors:** Xiaoyan Wei, Matthias Lienhard, Arunima Murgai, Julia Franke, Sophie Pöhle-Kronawitter, Georgios Kotsaris, Hao Wu, Stefan Börno, Bernd Timmermann, Rainer Glauben, Sigmar Stricker

## Abstract

Patients affected by neurofibromatosis type 1 (NF1) frequently show muscle weakness with unknown etiology. Here we show that Neurofibromin-1 (Nf1) is not required in muscle fibers, but specifically in early postnatal myogenic progenitors (MPs), where Nf1 loss led to cell cycle exit and differentiation blockade, depleting the MP pool resulting in reduced myonuclear accrual as well as reduced muscle stem cell numbers. This was caused by precocious induction of stem cell quiescence coupled to metabolic reprogramming of MPs impinging on glycolytic shutdown, which was conserved in muscle fibers. We show that a Mek/Erk/NOS pathway hypersensitizes Nf1-deficient MPs to Notch signaling, consequently, early postnatal Notch pathway inhibition ameliorated premature quiescence, metabolic reprogramming and muscle growth. This reveals an unexpected role of Ras/Mek/Erk signaling supporting postnatal MP quiescence in concert with Notch signaling, which is controlled by Nf1 safeguarding coordinated muscle growth and muscle stem cell pool establishment. Furthermore, our data suggest transmission of metabolic reprogramming across cellular differentiation, affecting fiber metabolism and function in NF1.

## Introduction

During prenatal development myogenic progenitors (MPs) that originate from the dermomyotome compartment of the somitic mesoderm migrate to their terminal destinations in the trunk and limbs. Here, MPs proliferate to expand the progenitor pool, from where cells continuously exit the cell cycle and differentiate into myoblasts under the control of myogenic regulatory factors Myf5, Mrf4 (Myf6), MyoD, and myogenin (Myog). Myoblasts ultimately fuse into multinucleated myofibers (Chal and Pourquie, 2017; Comai and Tajbakhsh, 2014). After birth, myofibers continue to grow by myonuclear accrual, carried by MPs that express the transcription factor Pax7. This process, however, gradually reduces and is replaced by metabolic fiber growth (White et al., 2010). Along with reduced myonuclear accrual, Pax7^+^ MPs exit the cell cycle and become quiescent. In parallel, they assume their characteristic position beneath the myofiber basal lamina; hence, they were called “satellite cells” (Mauro, 1961; Schultz et al., 1978). Adult satellite cells are muscle stem cells (MuSCs) that remain quiescent until activation by, for example, acute injury, after which they re-enter the cell cycle to establish a new MP pool to regenerate myofibers (Relaix and Zammit, 2012). In mice, most Pax7^+^ juvenile MPs enter quiescence by postnatal day 21 (p21) (Bachman et al., 2018; White et al., 2010). However, few proliferative Pax7^+^ MPs are still present until approx. p56 (Gattazzo et al., 2020) and even after, contributing to low, steady-state muscle turnover (Keefe et al., 2015; Pawlikowski et al., 2015). Quiescence of adult MuSC is maintained by a combination of niche signals, disruption of which leads to MuSC activation and cell cycle entry (Abou-Khalil et al., 2009; Baghdadi et al., 2018; Bjornson et al., 2012; Goel et al., 2017; Lukjanenko et al., 2019; Mourikis et al., 2012b; Wen et al., 2012). The primal establishment of MuSC quiescence during early postnatal life, however, is less understood. Notch signaling has been involved, as genetic perturbation disrupts MPs’ homing to the MuSC niche (Brohl et al., 2012) and disturbs the cell cycle exit of juvenile MPs and their transmission to quiescence (Al Jaam et al., 2016; Fukada et al., 2011; Kim et al., 2016).

*Neurofibromin 1* (*Nf1*) is a tumor suppressor gene that encodes a RAS–guanosine triphosphatase–activating protein (RAS-GAP) that reduces RAS activity and inhibits mitogen-activated protein kinase kinase 1/2–extracellular signal–regulated protein kinase 1/2 (Mek1/2-Erk1/2) downstream signaling (Ratner and Miller, 2015). Autosomal dominant *NF1* mutations cause neurofibromatosis type 1 (NF1), one of the most common genetic diseases affecting approx. 1 in 3000-4000 live births, which is mainly characterized by the presence of nerve sheath tumors (Ratner and Miller, 2015), in addition to musculoskeletal involvement, strongly affecting the patients’ mobility and quality of life (Elefteriou et al., 2009). NF1 patients, as well as those with mutations in other members of the RAS pathway (so-called RASopathies), often display severe muscle weakness (Souza et al., 2009; Stevenson et al., 2012; Summers et al., 2015). The function of *Nf1* in muscle is unclear. Constitutive inactivation of *Nf1* causes early embryonic lethality (Brannan et al., 1994), while *Nf1* haploinsufficiency does not affect muscle development of function in mice (Sullivan et al., 2014). Widespread *Nf1* inactivation in limb mesenchyme reduces muscle size and function (Kossler et al., 2011). Muscle-specific *Nf1* inactivation via Myod^Cre^ causes early postnatal lethality (Sullivan et al., 2014), while *Nf1* inactivation via Myf5^Cre^ causes viable offspring with muscle hypotrophy, fast fiber atrophy and a whole-body catabolic phenotype (Wei et al., 2020). Intriguingly, all three models indicated aberrant muscle metabolism with distorted carbohydrate and lipid usage (Sullivan et al., 2014; Summers et al., 2018; Wei et al., 2020).

In this study we show that *Nf1* is required in murine juvenile MPs to prevent metabolic reprogramming and precocious induction of stem cell quiescence. Loss of *Nf1* amplifies a Ras-Mek1/2-Erk1/2 signaling axis that funnels into Notch signaling. This distorts the balance between MP amplification/differentiation vs. quiescence induction, draining the postnatal MP pool, thus affecting postnatal muscle growth and MuSC pool establishment. Importantly, Nf1 was dispensable in muscle fibers, indicating that metabolic reprogramming of MPs can be transmitted to adult myofibers, and that the NF1-associated myopathy is a postnatal developmental disease.

## Results

### Premature cell cycle exit and impaired myogenic differentiation of Nf1^Myf5^ MPs reduces myonuclear accrual and MuSC numbers

Mice with conditional inactivation of *Nf1* targeted to myoblasts by using Myf5^Cre^ (Myf5^Cre^;Nf1^flox/flox^, or “Nf1^Myf5^”) reduce mTORC1-dependent anabolic myofiber growth during postnatal development in a variety of fore- and hindlimb muscles, already visible during the first 3 weeks of postnatal life (Wei et al., 2020). During juvenile development, however, muscle growth occurs by a combination of myonuclear cell accrual and metabolic growth (Bachman et al., 2018; Gattazzo et al., 2020; White et al., 2010), so we analyzed Pax7^+^ juvenile MP behavior in Nf1^Myf5^ mice. Reverse transcription–quantitative polymerase chain reaction (RT-qPCR) and RNA-sequencing (RNA-Seq) confirmed efficient decrease in *Nf1* messenger RNA (mRNA) in fluorescence-activated cell sorting (FACS)-isolated MPs at postnatal day 7 (p7) (Supplementary Fig. 1A). Myf5^Cre^ mice are haplo-insufficient for *Myf5*; however, *Myf5* expression was not affected in p7 MPs of Myf5^Cre/+^ mice compared to wild type controls (Supplementary Fig. 1B).

Proliferation rates of Nf1^Myf5^ Pax7^+^ cells assessed by Ki67 immunolabeling of tissue sections were slightly reduced at p7, with a high decrease at p14 (Fig. 1A). At p21, Pax7^+^ cells appeared mostly non-proliferative in Nf1^Myf5^ muscles (Fig. 1A, B). A low fraction of adult MuSCs is in the cell cycle (Chakkalakal et al., 2012), which was also seen at p84 in controls but not in Nf1^Myf5^ Pax7^+^ MuSCs (Fig. 1A). Freshly FACS-isolated p14 MPs after cytospin confirmed a reduced proliferation index (Fig. 1C). In addition, p14 Nf1^Myf5^ MPs showed increased Pax7 protein abundance, indicated by fluorescence intensity measurement (Fig. 1C).

**Fig. 1.**
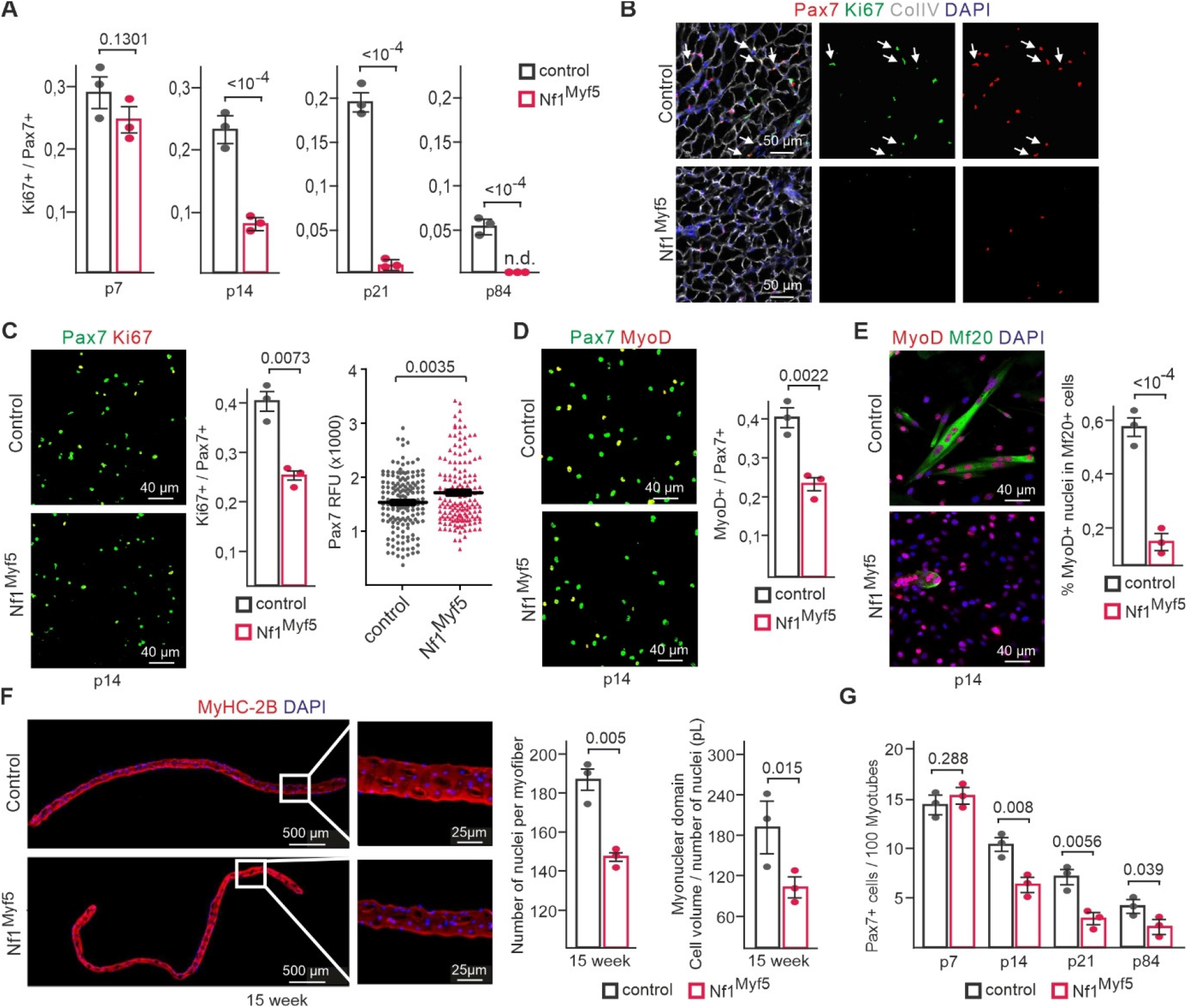
Premature cell cycle exit and impaired differentiation of Nf1^Myf5^ MPs reduces myonuclear accrual and MuSC numbers. **A)** Ki67^+^/Pax7^+^ cells quantification relative to all Pax7^+^ cells in TA muscles of control and Nf1^Myf5^ mice at indicated time points. p, postnatal day (*n* = 3). **B)** Representative immunolabeling images of Pax7 and Ki67 of p21 muscle sections of control or Nf1^Myf5^ mice. Arrows indicate Pax7^+^/Ki67^+^ cells (*n* = 3). **C)** Cytospin of FACS-isolated p14 MPs from control or Nf1^Myf5^ mice labeled for Pax7 and Ki67. Quantification of Ki67^+^/Pax7^+^ cells and Pax7 relative fluorescence intensity (RFU) (right) (*n* = 3; >200 cells/sample). **D)** Cytospin of FACS-isolated p14 MPs from control or Nf1^Myf5^ mice labeled for Pax7 and MyoD. Quantification of Pax7^+^/MyoD^+^ cells (right) (*n* = 3; >200 cells/sample). **E)** *In vitro* differentiation of FACS-isolated p14 MPs from control or Nf1^Myf5^ mice. (Left) Representative images after 2 d differentiation stained for Myosin (Mf20) and MyoD. (Right) Quantification of MyoD^+^ nuclei within myofibers relative to all MyoD^+^ nuclei (*n* = 3; >200 cells/sample). **F)** Top: Representative images of single fibers isolated from 15-week EDL muscles stained for MyHC-2B (red) and DAPI (nuclei, blue). Boxed region shown as magnification. Below: Quantification of nuclei per myofiber; pL, picoliter, and myonuclear domain (cell volume/number of nuclei) (*n* = 3; >30 fibers/sample). **G)** Pax7^+^ cell quantification in TA muscles of control or Nf1^Myf5^ mice at indicated time points (*n* = 3). Data are mean ± SEM; *P*-value calculated by two-sided unpaired *t*-test; *N*-numbers indicate biological replicates (mice per genotype).

Cytospun p14 MPs showed a decrease in MyoD^+^/Pax7^+^ cell numbers, indicating a differentiation delay (Fig. 1D), which was confirmed by plating freshly isolated p14 MPs in high density and immediately subjecting them to differentiation conditions; we saw a strong decrease in myotube formation from Nf1^Myf5^ MPs (Fig. 1E). These results indicated cell cycle exit of Nf1^Myf5^ MPs that was not coupled to myoblast differentiation.

Decreased proliferation and differentiation reduced myonuclear accrual in Nf1^Myf5^ mice, as shown by reduced myonuclear numbers in single fast (MyHC-2B^+^) fibers from extensor digitorum longus (EDL) muscles of adult (15-week-old) mice (Fig. 1F). In addition, we found a reduced myonuclear domain (the amount of cytoplasm allocated to one myonucleus) in Nf1^Myf5^ fibers (Fig. 1F), suggesting no compensatory myonuclear domain growth (Cramer et al., 2020) occurred in this model, consistent with the metabolic growth defect of Nf1^Myf5^ muscle (Wei et al., 2020). Pax7^+^ cell numbers of Nf1^Myf5^ muscles were normal at p7 but reduced in the following 2 weeks of postnatal life, and at p21, Pax7^+^ cell numbers in Nf1^Myf5^ muscles reduced to ∼50% of control levels (Fig. 1G) and remained constant thereafter, as found at p84 (Fig. 1G), indicating a lasting decrease in MuSC numbers. We did not detect aberrant apoptosis in Nf1^Myf5^ muscle (Supplementary Fig. 1C).

We conclude that precocious postnatal cell cycle withdrawal of Pax7^+^ MPs and a differentiation blockade explain the decrease in MP numbers and myonuclear accretion, as well as the diminished adult MuSC pool in *Nf1* mutants.

### Nf1 is dispensable in muscle fibers

Myf5^Cre^ targets myogenic progenitors (myoblasts), thus leading to an early recombination in the majority of the myogenic lineage (Comai et al., 2014). To disentangle the function of *Nf1* within myofibers uncoupled from an earlier function in MPs, we inactivated *Nf1* using Acta1^Cre^, which targets myofibers but not myoblasts (Miniou et al., 1999). Acta1^Cre^ specificity in limb muscle fibers, but not Pax7^+^ myogenic progenitors (MPs), was confirmed in Rosa26^mTmG^ reporter mice (Muzumdar et al., 2007) (Supplementary Fig. 2A, B). Surprisingly. Acta1^Cre^;Nf1^flox/flox^ mice (Nf1^Acta1^) showed normal growth and were indistinguishable from littermates (Fig. 2A).

**Fig. 2.**
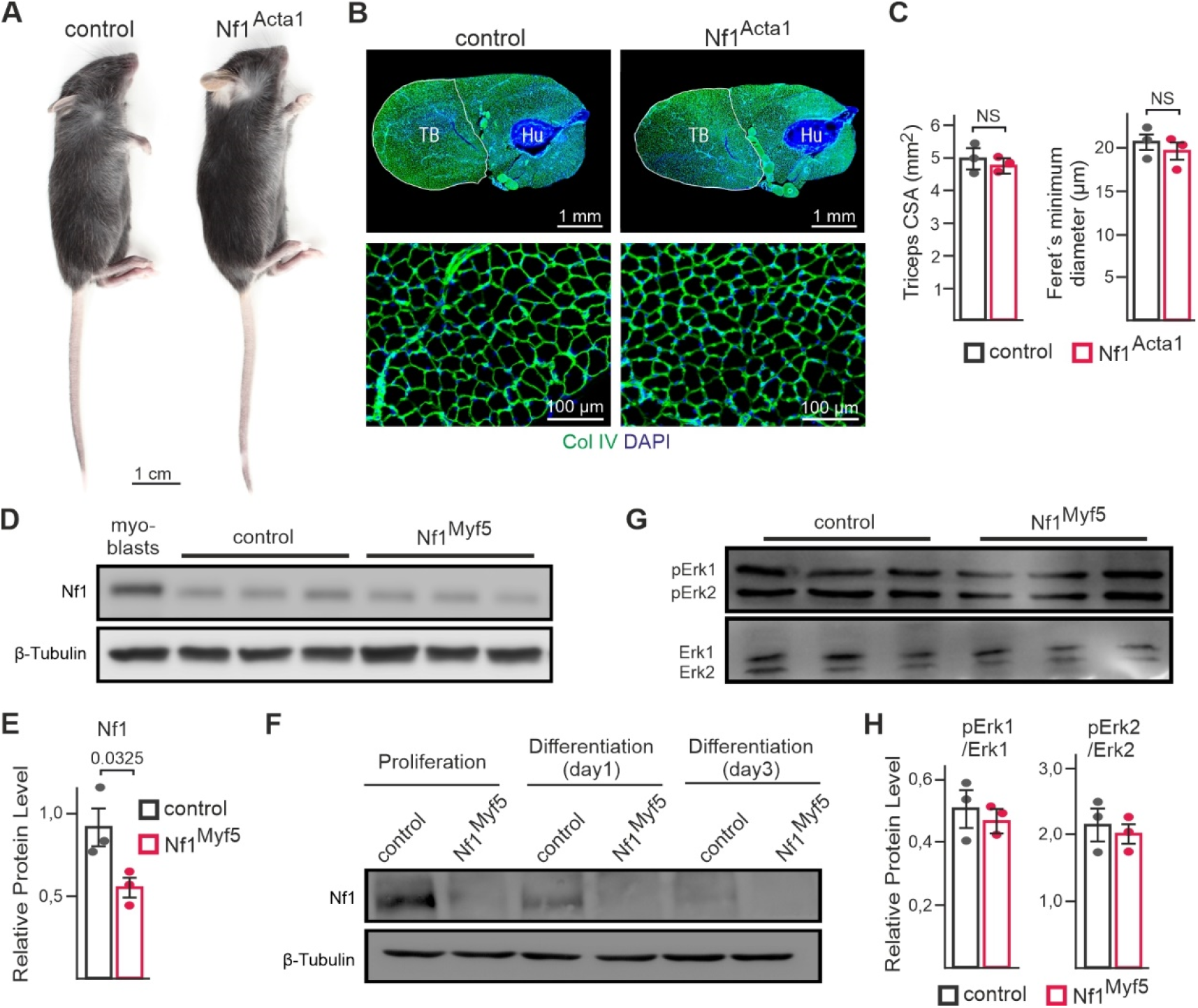
Nf1 is dispensable in myofibers. **A)** Whole-body appearance of control and Nf1^Acta1^ mice at p21. **B)** Cross sections through upper forelimbs of control and Nf1^Acta1^ mice at p21. TB, triceps brachii; Hu, humerus. Magnification from TB muscle shown below. **C)** Quantification of TB cross-sectional area (left) and myofiber feret’s diameter (right) of control and Nf1^Acta1^ TB muscle (*n* = 3). **D)** Western blot analysis of Nf1 protein expression in p21 muscle tissue; p21 wild type primary myoblasts are shown for comparison. **E)** Quantification of (D) (*n*=3). **F)** Western blot analysis of Nf1 expression in control and Nf1Myf5 primary myoblasts in proliferation medium, and after 1 or 3 days in differentiation medium. **G)** Western blot analysis of Erk1/2 phosphorylation in p21 muscle tissue of control and Nf1Myf5 mice. **H**) quantification of (G) (n=3). Data are mean ± SEM; *P*-value calculated by two-sided unpaired *t*-test. N-numbers indicate biological replicates (mice per genotype).

The whole triceps muscle cross-sectional area and fiber diameters were equal to controls (Fig. 2B, C). RT-qPCR confirmed the efficiency of *Nf1* deletion in p21 muscle tissue (Supplementary Fig. 2C).

We therefore analyzed Nf1 protein expression and Erk1/2 pathway activity during myogenic differentiation. Nf1 protein was barely detectable by western blot in p21 muscle lysate, as opposed to marked expression in primary myoblasts (Fig. 2D). Nevertheless, a downregulation of Nf1 protein in Nf1^Myf5^ samples was detected (Fig. 2E). Primary myoblast culture showed that Nf1 expression is downregulated early during myogenic differentiation (Fig. 2F). Finally, muscle protein lysates from Nf1^Myf5^ mice did not show a difference in Erk1/2 phosphorylation compared to controls (Fig. 2G, H).

We conclude that Nf1 is not required in mature muscle fibers, suggesting that myopathy of Nf1^Myf5^ animals was caused by aberrant progenitor behavior.

### Nf1-deficient MPs are shifted to premature quiescence

Cell cycle exit with a lack of differentiation and increased Pax7 expression indicates a shift of MPs to a quiescent phenotype. To further address this at the phenotype onset, we analyzed freshly FACS-isolated p7 MPs, immediately fixed after isolation to preserve the *in vivo* condition. First, we analyzed freshly isolated Nf1^Myf5^ and control p7 MPs by RNA-Seq (Fig. 3A and Supplementary Fig. 3A, B). Gene set enrichment analysis (GSEA) confirmed upregulation of the RAS signaling pathway in Nf1^Myf5^ MPs (Fig. 3B), and p-Erk immunolabeling intensity increased in cytospun Nf1^Myf5^ MPs (Fig. 3C). Consistent with the reduced differentiation potential of Nf1^Myf5^ MPs, myogenesis-related GSEA terms were enriched in controls (Fig. 3D). In contrast, GSEA terms associated with the ECM and basal lamina, both essential for MuSC quiescence (Baghdadi et al., 2018; Yin et al., 2013), were overrepresented in mutants (Supplementary Fig. 3C). Kyoto Encyclopedia of Genes and Genomes (KEGG) analysis of differentially regulated genes confirmed “ECM–receptor interaction” and “Focal adhesion” among the highest-enriched terms in genes upregulated in Nf1^Myf5^ MPs (Supplementary Fig. 3D).

**Fig. 3.**
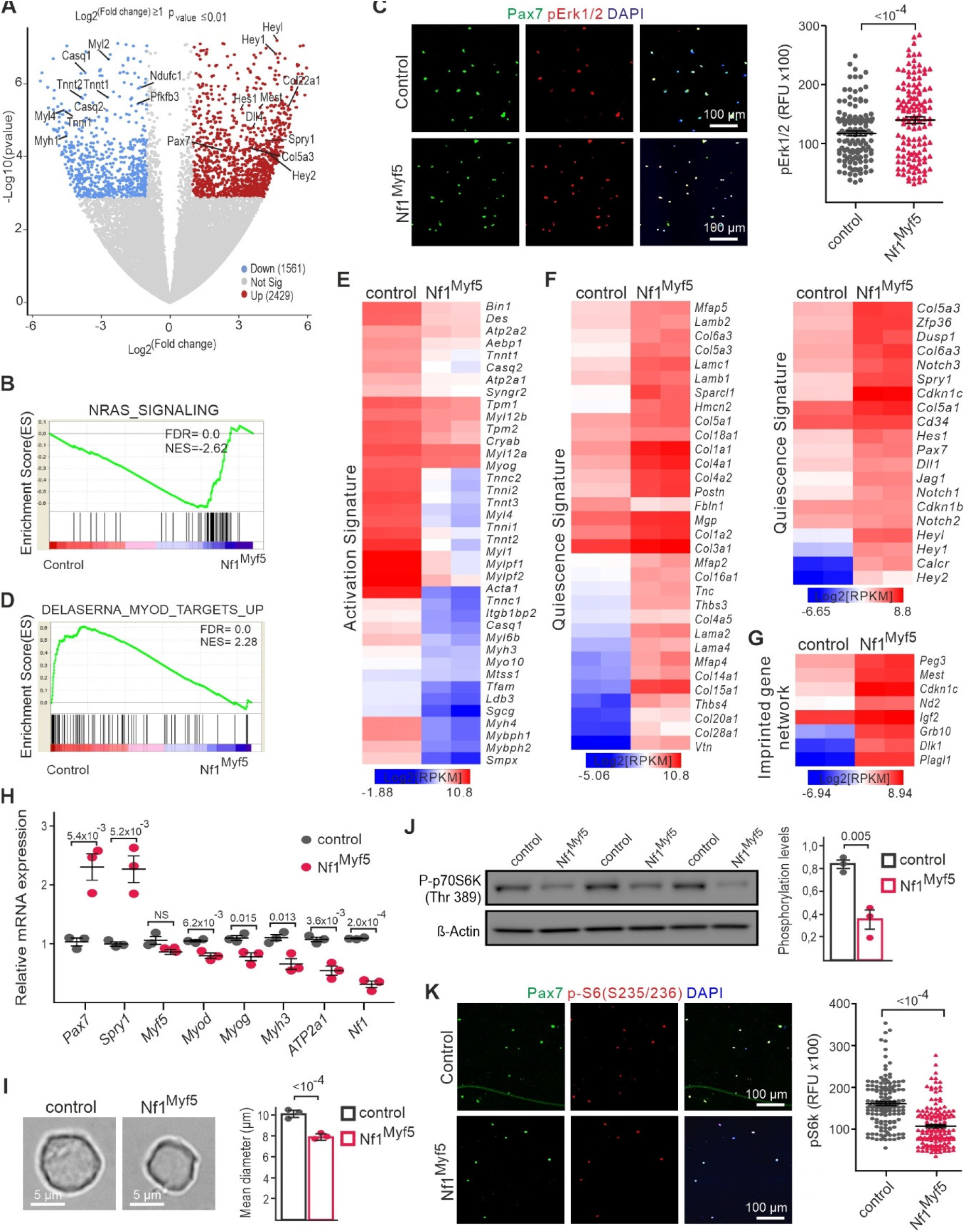
Premature shift of Nf1^Myf5^ MPs to quiescence. **A)** Volcano plot of transcriptome data from freshly FACS-isolated control or Nf1^Myf5^ p7 MPs. Individual transcripts deregulated in Nf1^Myf5^ MPs are indicated (blue: down; red: up). **B)** GSEA of control and Nf1^Myf5^ p7 MP RNA-Seq data for “NRAS Signaling”. **C)** GSEA of RNA-Seq data from control or Nf1^Myf5^ p7 MPs shows “MyoD targets” enriched in controls. **D)** Quantification of fluorescence signal intensity for pErk1/2 in control and Nf1^Myf5^ p7 Pax7^+^ MPs (n=3 animals per genotype). **E)** Heatmap shows globally reduced MuSC activation–related gene expression in p7 Nf1^Myf5^ MPs. **F)** Heatmap shows globally increased MuSC quiescence-related gene expression in p7 Nf1^Myf5^ MPs. **G)** Heatmap shows increased expression of imprinted gene network genes in p7 Nf1^Myf5^ MPs. **H)** RT-qPCR confirmation of differential expression of indicated genes in Nf1^Myf5^ p7 MPs (*n* = 3; each dot represents the mean of three technical replicates from one biological replicate). **I)** Reduced cell diameter in p7 Nf1^Myf5^ freshly sorted MPs. Representative images (left); quantification (right) (*n* = 3; ≥200 cells measured from each mouse). **J)** Western blot shows reduced p70s6 kinase phosphorylation at Thr-389 in p7 Nf1^Myf5^ MPs (*n* = 3). **K)** Quantification of fluorescence signal intensity for Serine-235/236-phosphorylated S6 ribosomal protein (p-S6) in control and Nf1^Myf5^ p7 MPs (n=3). Data are mean ± SEM; *P*-value calculated by two-sided unpaired *t*-test; *N*-numbers indicate biological replicates (mice per genotype).

We compared differentially expressed genes (DEGs) to MuSC activation and quiescence signatures, obtained from published transcriptome datasets (Baghdadi et al., 2018; Machado et al., 2017; Pallafacchina et al., 2010; Ryall et al., 2015; Tierney et al., 2016; van Velthoven et al., 2017) mined for genes commonly down- or upregulated. MuSC activation–related genes, including *Myog*, were downregulated in Nf1^Myf5^ MPs (Fig. 3E), while MuSC quiescence-related genes were uniformly upregulated in Nf1^Myf5^ MPs (Fig. 3F). This included *Cdkn1b* and *Cdkn1c*, which encode cell cycle inhibitors p27 and p57, consistent with cell cycle exit. The so-called imprinted gene network, known to be highly expressed in quiescent stem cells (Berg et al., 2011; Besson et al., 2011), was upregulated in Nf1^Myf5^ MPs (Fig. 3G). Global comparison of RNA-Seq data to that of Ryall et al. (2015) confirmed a shift of Nf1^Myf5^ MPs transcriptome toward the signature of quiescent MuSCs (Supplementary Fig. 3E).

RT-qPCR confirmed upregulation of quiescence-related genes *Pax7* and *Spry1*, and downregulation of activation–related genes *Myod, Myog*, and *Myh3* and *ATP2a1* (Fig. 3H). Freshly isolated Nf1^Myf5^ MPs were smaller than control cells (Fig. 3I), a known feature of quiescent MuSCs (Rodgers et al., 2014). In addition, phosphorylation of p70S6 kinase and of S6 ribosomal protein serine-235/236 as mammalian target of rapamycin complex 1 (mTORC1) signaling readouts, which is known to be induced upon MuSC activation (Rodgers et al., 2014), was reduced in Nf1^Myf5^ p7 MPs (Fig. 2J, K).

Results indicated that Nf1^Myf5^ p7 MPs show a phenotypic shift toward MuSC quiescence, including a transcriptomic signature, cell cycle exit, and mTORC1 activity.

### Quiescence shift of Nf1^Myf5^ MPs reflects an altered epigenetic landscape overlapping that of adult quiescent MuSCs

GSEA analysis of transcriptome data showed overrepresentation of “condensed chromosome”-related genes to be upregulated in p7 Nf1^Myf5^ MPs (Fig. 4A), indicating chromatin alterations. We assessed three major chromatin marks: histone 3 lysine 4 trimethylation (H3K4me3), generally associated with active promoters; H3K27me3, associated with transcription inhibition (Kouzarides, 2007); and DNA methylation (5’cytosine), globally associated with transcription inhibition (Greenberg and Bourc’his, 2019). Chromatin immunoprecipitation sequencing (ChIP-Seq) analysis of freshly isolated p7 MPs showed apparently unchanged H3K4me3 levels between controls and Nf1^Myf5^ MPs (Fig. 4B), while H3K27me3 levels globally reduced in Nf1^Myf5^ MPs (Fig. 4C), as confirmed by immunolabeling (Supplementary Fig. 4A). We observed a decrease in H3K27me3 decoration at quiescence-related genes, such as *Pax7* (Fig. 4D), or cell cycle repression genes, such as *Cdkn2a* (Supplementary Fig. 4B). Therefore, acquisition of a quiescent state in Nf1^Myf5^ MPs is concomitant with reduced repressive H3K27me3 modification at quiescence-related genes. RNA-Seq analysis showed upregulation of DNA-demethylase *Tet1-3* and all three relevant DNA methyltransferases, *Dnmt1, Dnmt3a*, and *Dnmt3b* (Fig. 4E). The deregulation of these genes overlapped the signature in quiescent MuSCs (Ryall et al., (2015) (Supplementary Fig. 4C). *Dnmt1* and *Dnmt3a* were highly expressed in p7 MPs, but *Dnmt3b* had low expression (Supplementary Data 1). RT-qPCR confirmed *Dnmt1* and *Dnmt3a* upregulation in Nf1^Myf5^ MPs (Fig. 4F). Methylated DNA immunoprecipitation sequencing (MeDIP-Seq) analysis of freshly isolated p7 MPs showed differential methylation between controls and Nf1^Myf5^ predominantly at CpG islands (Fig. 4G). Most differentially methylated CpG islands were hypermethylated in Nf1^Myf5^ MPs (Fig. 4G).

**Fig. 4.**
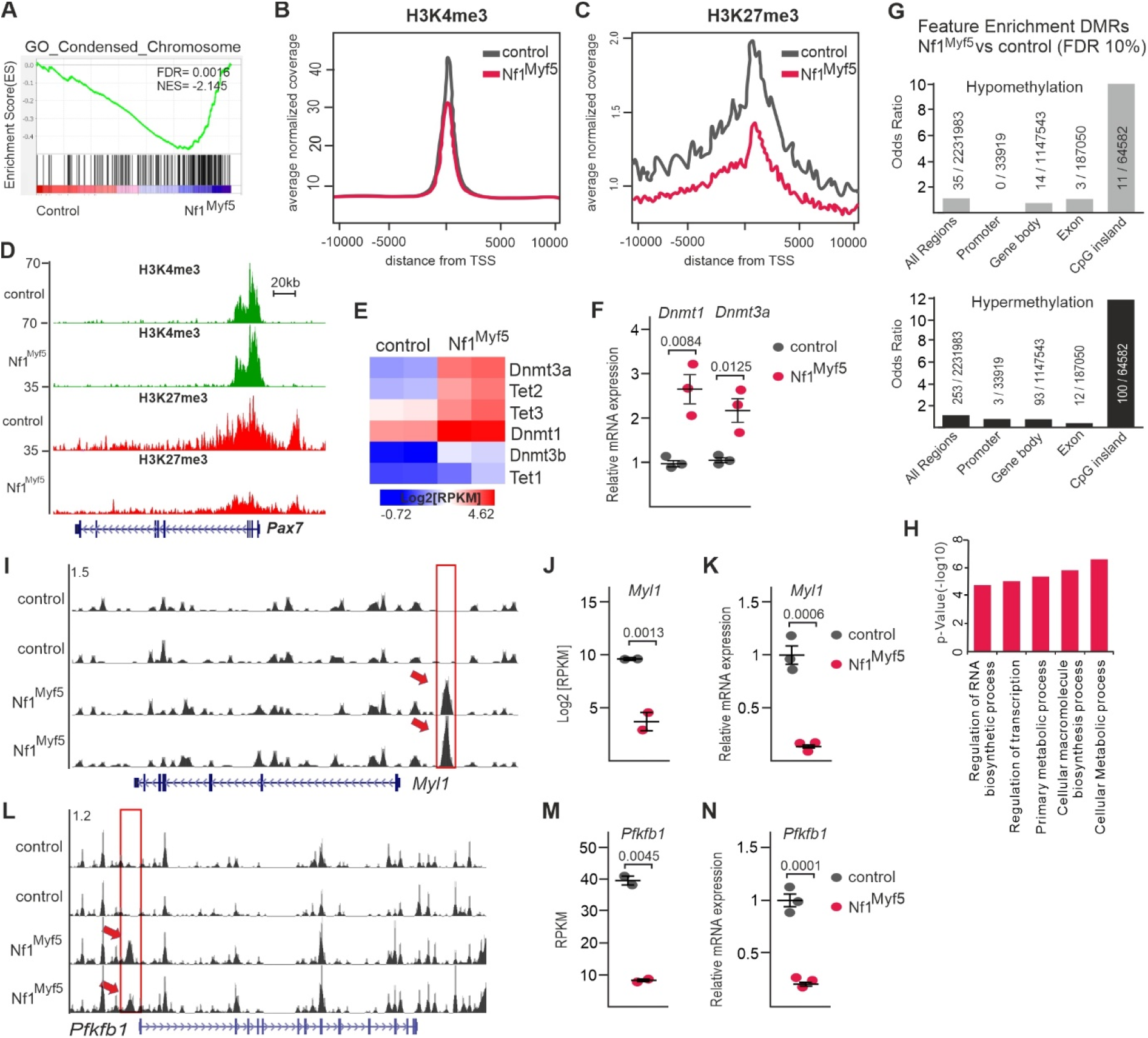
Epigenetic changes associated with quiescence shift of Nf1^Myf5^ MPs. **A)** GSEA of control and Nf1^Myf5^ p7 MP RNA-Seq data for “condensed chromosome.” **B, C)** Averaged normalized coverage for H3K4me3 and H3K27me3, derived from ChIP-Seq performed on control and Nf1^Myf5^ FACS-isolated p7 MPs. TSS, transcription start site. **D)** ChIP-Seq tracks for H3K4me3 and H3K27me3 at the *Pax7* locus in control and Nf1^Myf5^ p7 MPs. **E)** Heatmap depiction of DNA methlaytion–related gene expression in control and Nf1^Myf5^ p7 MPs. **F)** RT-qPCR of *Dnmt1* and *Dnmt3a* in control and Nf1^Myf5^ p7 MPs (*n* = 3). **G)** Enrichment of hypo- and hypermethylated regions among differentially methylated regions (DMRs) in control and Nf1^Myf5^ p7 MPs. **H)** GO analysis of hypermethylated DMRs in Nf1^Myf5^ p7 MPs. **I)** MeDIP-Seq tracks from control and Nf1^Myf5^ p7 MPs at the *Myl1* locus. **J)** Log2(RPKM) values for *Myl1* in p7 MPs transcriptome data (*n* = 2). **K)** RT-qPCR of *Myl1* expression in control and Nf1^Myf5^ p7 MPs (*n* = 3). **L)** MeDIP-Seq tracks from control and Nf1^Myf5^ p7 MPs at the *Pfkfb1* locus. **M)** RPKM values for *Pfkfb1* in p7 MP transcriptome data (*n* = 2). **N)** RT-qPCR of *Pfkfb1* expression in control and Nf1^Myf5^ p7 MPs (*n* = 3). Data are mean ± SEM; *P*-value calculated by two-sided unpaired *t*-test; *N*-numbers indicate biological replicates (mice per genotype).

Gene Ontology (GO) overrepresentation analysis of genes in proximity to hypermethylated CpG islands showed terms associated with RNA synthesis and transcription, and cellular metabolism (Fig. 4H). Mining the proximity of differentially methylated regions for myogenesis-related genes showed a hypermethylated CpG island 3.5 kilobases (kb) upstream of *Myl1* (Fig. 4I), which is part of the activation signature and is downregulated in Nf1^Myf5^ MPs in transcriptome data (Fig. 4J). RT-qPCR confirmed *Myl1* downregulation (Fig. 4K).

Analysis of metabolism-related genes showed hypermethylation of a CpG island 1.5 kb upstream of *Pfkfb1*, encoding phosphofructo-kinase-fructose-bisphosphatase 1 (Fig. 4L). *Pfkfb1* was the highest expressed isoform of all *Pfkfb*s in juvenile MPs (Supplementary Data 1), and *Pfkfb1* mRNA expression was strongly downregulated in Nf1^Myf5^ MPs in transcriptome data (Fig. 4M) and RT-qPCR (Fig. 4N). In addition, hypermethylation overlapped the promoter of *Ndufb11*, encoding a subunit of mitochondrial complex I, and *Idh3g*, encoding a subunit of mitochondrial isocitrate dehydrogenase, which catalyzes the rate-limiting step of the tricarboxic acid cycle (TCA) (Supplementary Fig. 4D).

Therefore, epigenetic alterations observed in Nf1^Myf5^ MPs are consistent with a shift toward quiescence. These alterations could contribute to impaired myogenic differentiation, and hypermethylation and transcriptional downregulation of metabolic genes indicate changes in cellular energy metabolism in Nf1^Myf5^ MPs.

### Metabolic reprogramming of Nf1^Myf5^ juvenile MPs that is conserved in myofibers

Consistent with possible metabolic alterations, overrepresentation analysis with KEGG database pathways of the p7 MP transcriptome showed enrichment of “metabolic pathways,” “carbon metabolism,” “biosynthesis of amino acids,” and “glycolysis/gluconeogenesis” in genes downregulated in Nf1^Myf5^ MPs, and “protein digestion and absorption” in upregulated genes (Supplementary Fig. 3D). GSEA showed enrichment of “glycolysis/gluconeogenesis” and “oxidative phosphorylation” in controls (Fig. 5A). We identified global downregulation of genes of the glycolytic pathway and the pyruvate dehydrogenase complex, the citrate cycle, and the mitochondrial electron transport chain (ETC) in Nf1^Myf5^ p7 MPs (Fig. 5B–D).

**Fig. 5.**
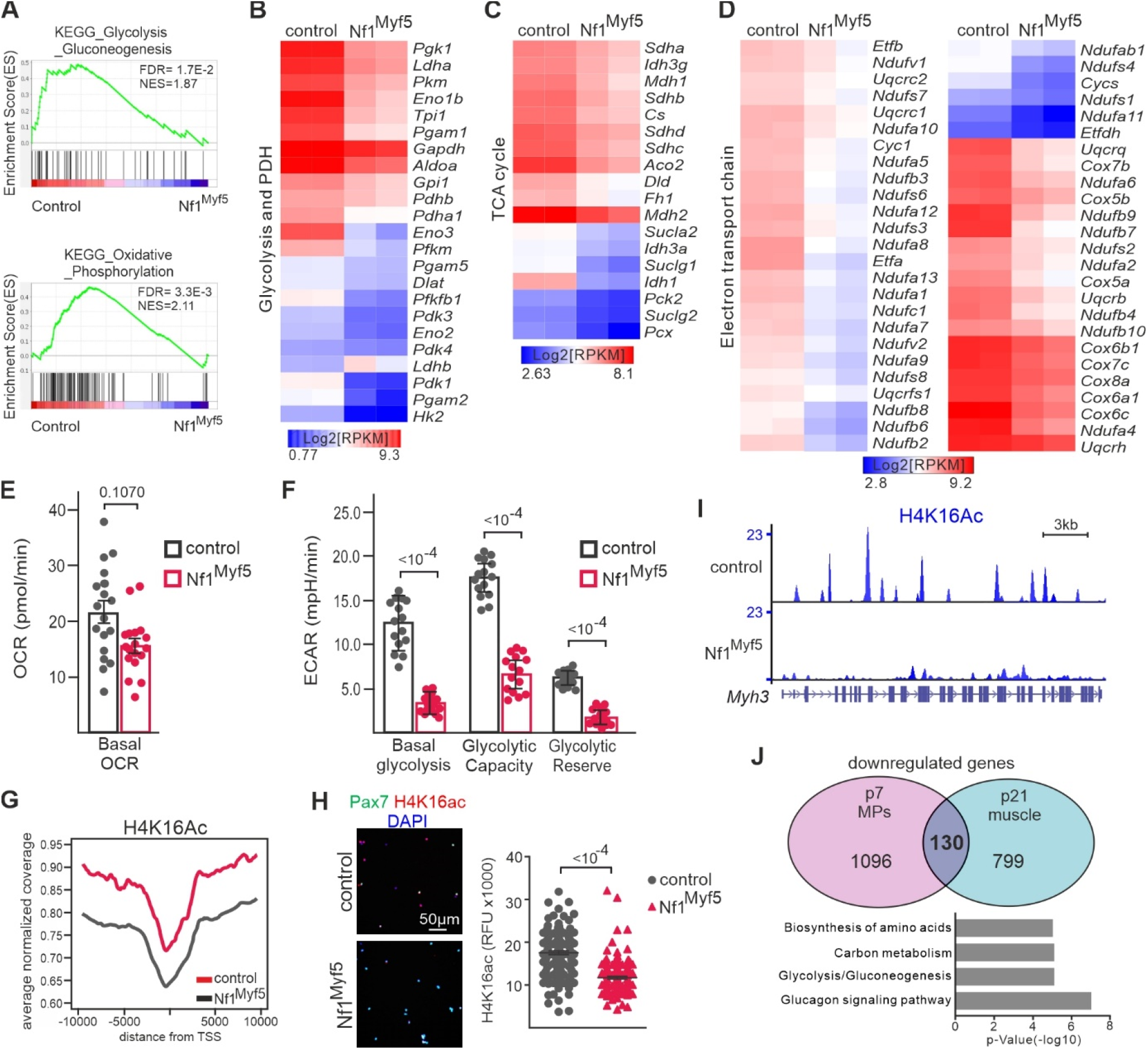
Metabolic reprogramming of Nf1^Myf5^ MPs. **A)** GSEA of control and Nf1^Myf5^ p7 MP RNA-Seq data for “glycolysis - gluconeogenesis” and “oxidative phosphorylation.” **B)** Heatmap shows glycolysis-related gene downregulation in Nf1^Myf5^ MPs. **C)** Heatmap shows TCA cycle component gene expression decrease in Nf1^Myf5^ MPs. **D)** Heatmap shows ETC component gene expression decrease in Nf1^Myf5^ MPs. **E)** SeahorseXF flux analysis of control and Nf1^Myf5^ p7 MPs; quantification of OCR (*n* = 3). **F)** SeahorseXF flux analysis of control and Nf1^Myf5^ p7 MPs; quantification of ECAR (*n* = 3). **G)** Averaged normalized coverage for H4K16 derived from ChIP-Seq on control and Nf1^Myf5^ p7 FACS-isolated MPs. TSS, transcription start site. **H)** Immunolabeling for H4K16ac on FACS isolated cytospun MPs from p7 control and Nf1^Myf5^ animals; quantification is shown right (n=3). **I)** ChIP-Seq tracks for H4K16ac from control and Nf1^Myf5^ p7 MPs at the *Myh3* locus. **J)** Venn diagram showing 130 commonly downregulated genes between Nf1^Myf5^ p7 MPs and Nf1^Myf5^ p21 muscle. GO analysis of commonly downregulated genes shown below. Data are mean ± SEM; *P*-value calculated by two-sided unpaired *t*-test; N-numbers indicate biological replicates (mice per genotype).

Seahorse metabolic flux analysis with freshly isolated p7 MPs to assess the metabolic consequence of this deregulation showed a moderate decrease in the basal oxygen consumption rate (OCR) below statistical significance in Nf1^Myf5^ p7 MPs, indicating that oxidative metabolism is only mildly affected (Fig. 5E). In contrast, the extracellular acidification rate (ECAR) was highly reduced in Nf1^Myf5^ MPs, indicating severe glycolytic flux inhibition (Fig. 5F). Therefore, Nf1^Myf5^ MPs mainly use low-level oxidative metabolism as an energy source, consistent with a quiescent phenotype (Ryall and Lynch, 2018).

A metabolic switch from slow oxidative to forced glycolytic metabolism occurs during adult MuSC exit from quiescence to activation (Ryall et al., 2015). Concomitant NAD^+^ depletion inhibits Sirt1 function, which acts as a histone deacetylase mainly targeting H4K16. Increased H4K16 acetylation upon muscle-specific Sirt1 deletion induces expression of MuSC activation and myogenic differentiation-related genes (Ryall et al., 2015). H4K16Ac ChIP-Seq showed a decrease in global levels of H4K16Ac in p7 Nf1^Myf5^ MPs (Fig. 5G). Analysis of global H4K16ac levels in freshly isolated p7 MPs (Fig. 5H) and in p7 and 12-week-old muscle sections by immunofluorescence (Supplementary Fig. 5A, B) confirmed long-term decreased H4K16ac levels in Nf1^Myf5^ Pax7^+^ cells compared to controls.

*Myh3, Bgn, Fst* and *Mylk2*, which are upregulated in MuSC-specific Sirt1 conditional mice (Ryall et al., 2015), were downregulated in Nf1^Myf5^ MPs (Supplementary Fig. 5C) and showed reduced H4K16Ac decoration at their gene bodies (Fig. 5I and Supplementary Fig. 5D-F). Thus, Nf1^Myf5^ MPs are driven toward quiescence and show metabolic reprogramming with severely inhibited glycolytic metabolism, and decreased H4K14ac decoration and expression of myogenic differentiation-related genes.

Differentiated muscle in Nf1^Myf5^ animals shows fast fiber atrophy concomitant with stunted glycolysis (Wei et al., 2020). Comparing transcriptome data of Nf1^Myf5^ p7 MPs to Nf1^Myf5^ p21 muscle (Wei et al., 2020) showed common downregulation of 130 genes (Fig. 5J). GO overrepresentation analysis of this gene set showed enrichment of terms related to glucose/carbon metabolism and amino acid synthesis (Fig. 5J). This overlap in Nf1-deficient progenitors and differentiated muscle indicates a continuous deregulation of carbohydrate metabolism and anabolic pathways in Nf1-deficient muscle. As Nf1^Acta1^ mice show no muscle defect, this suggests that perturbed muscle fiber metabolism in Nf1^Myf5^ animals (Wei et al., 2020) can be traced back to defects in juvenile MPs, suggesting that that metabolic reprogramming in Nf1^Myf5^ juvenile MPs is transmitted to myofibers.

### Increased Notch signaling induced by a Mek/Erk/NOS cascade drives Nf1^Myf5^ juvenile MP quiescence shift

To analyze the mechanism of Nf1^Myf5^ MPs quiescence shift and reprogramming, we performed *in vitro* culture of juvenile primary myoblasts. RT-qPCR confirmed effective *Nf1* knockdown (Supplementary Fig. 6A). Surprisingly, *in vitro*, Nf1^Myf5^ myoblasts did not reproduce the *in vivo* proliferative behavior, but showed enhanced proliferation (Supplementary Fig. 6B). Switching cells to a differentiation medium after 2 d of culture showed effective block of myogenic differentiation in Nf1^Myf5^ primary MPs (Supplementary Fig. 6B), as observed before for FACS-isolated MPs. Both increased proliferation and blocked differentiation fully depended on Mek/Erk signaling, as shown by inhibition with UO126 (Supplementary Fig. 6B). The discrepancy between *in vivo* and *in vitro* proliferation behavior of Nf1^Myf5^ MPs indicated that *in vivo* non-cell-autonomous microenvironmental factors override or divert Mek/Erk signaling in juvenile MPs, inhibiting cell proliferation.

Transcriptome analysis indicated upregulation of Delta/Notch signaling pathway components *Dll1* and *Notch1/2*, and upregulation of Notch pathway targets *Hes1, Hey1, Heyl, Calcr*, and *Col5a1* (Fig. 3F). GSEA showed enrichment of “Notch targets” in Nf1^Myf5^ MPs (Supplementary Fig. 6C). Upregulation of *Notch1, Notch3, Hes1*, and *Hey1* in p7 Nf1^Myf5^ MPs was confirmed by RT-qPCR (Fig. 6A). We therefore tested whether Notch signaling represents the *in vivo* niche factor lacking *in vitro* by culturing MPs on cell culture plates coated with recombinant Jagged-1, which activates Notch signaling in myogenic cells (Mu et al., 2015; Safaee et al., 2017; Sakai et al., 2017).

**Fig. 6.**
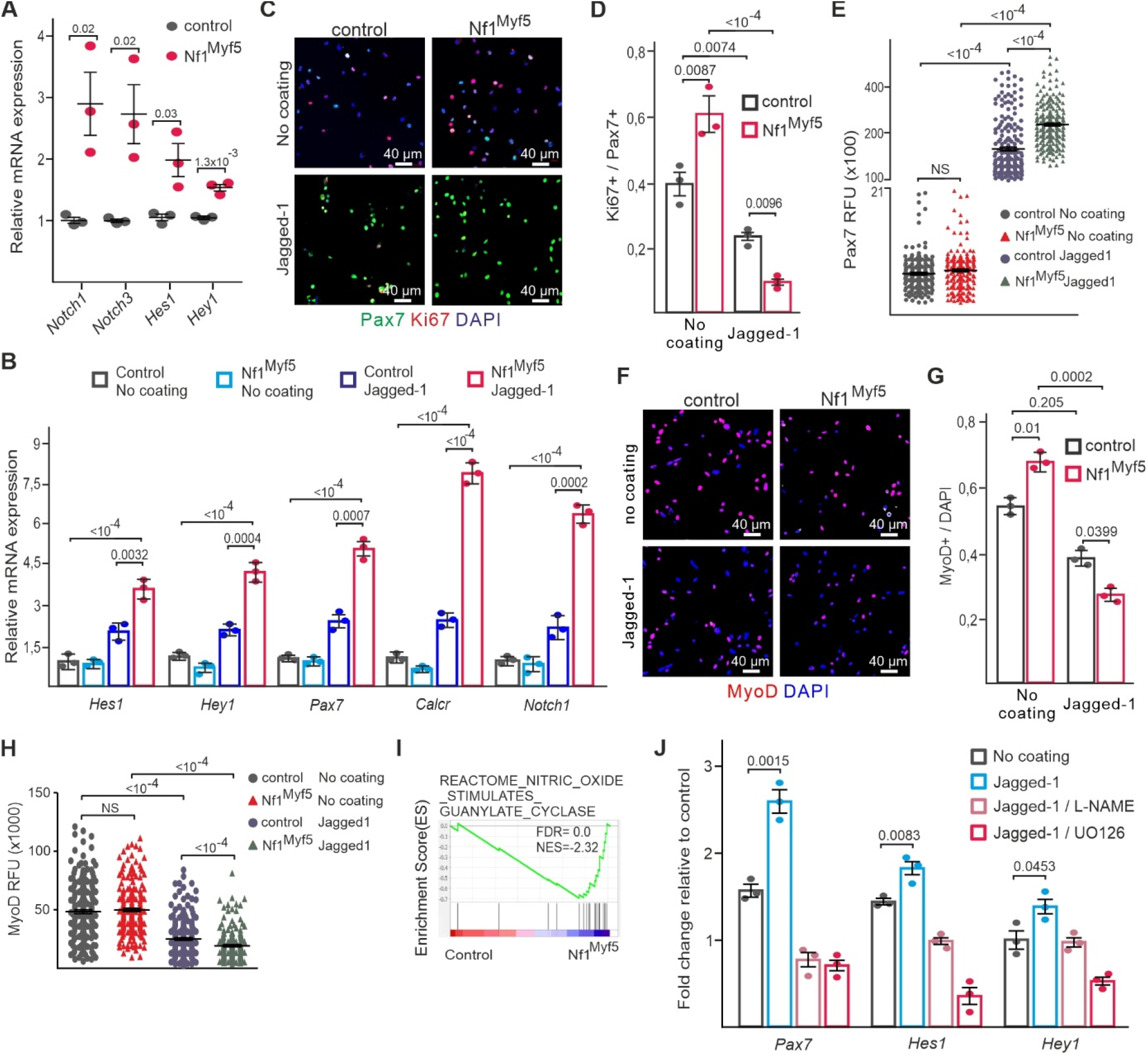
Increased Notch signaling drives Nf1^Myf5^ MPs to quiescence. **A)** RT-qPCR of Notch pathway component and target genes in control and Nf1^Myf5^ p7 MPs (*n* = 3; each dot represents the mean of three technical replicates from one biological replicate). **B)** RT-qPCR for Notch targets on FACS-isolated control or Nf1^Myf5^ p14 MPs cultured on Matrigel without coating or Jagged-1 coating for 48 h (*n* = 3). **C)** FACS-isolated control or Nf1^Myf5^ p14 MPs cultured on Matrigel without coating or Jagged-1 coating for 48 h stained for Pax7 and Ki67. **D)** Ki67^+^ cell quantification among Pax7^+^ cells on image data as in (B) (*n* = 3). **E**) Pax7 fluorescence intensity quantification on image data as in (B) (*n* = 3). **F)** Immunolabeling for MyoD on FACS-isolated control or Nf1^Myf5^ p14 MPs cultured on Matrigel w/o coating or Jagged-1 coating for 48 hours. **G)** Quantification of numbers of MyoD^+^ cells / total cells on image data as in (F) (*n* = 3). **H)** Quantification of MyoD fluorescence intensity on image data as in (B) (*n* = 3). **I)** GSEA on RNA-Seq data from control and Nf1^Myf5^ p7 MPs for “nitric oxide stimulates guanylate cyclase”. **J)** FACS-isolated control or Nf1^Myf5^ p14 MPs cultured on Matrigel without coating or Jagged-1 coating for 48 h with or without Mek inhibitor UO126 or pan-NOS inhibitor L-NAME. RT-qPCR for Notch targets performed. Bars show fold-changes of Nf1^Myf5^ MPs relative to control MPs, set as 1 (*n* = 3). Data are mean± SEM; *P*-value calculated by two-sided unpaired *t*-test; *N*-numbers indicate biological replicates (mice per genotype).

To calibrate the system, we first cultured wild type primary myoblasts for 2 d in proliferation medium on uncoated control dishes, or dishes coated with different concentrations of Jagged-1 ligand. This showed induction of *Hes1* and *Hey1* expression already at 2,5 ng/μl, with maximal induction reached at 5 ng/μl (Supplementary Fig. 6D). Compared to control cells, Nf1^Myf5^ primary myoblasts showed increased induction of Notch target gene expression on recombinant Jagged-1–coated (5 ng/μl) dishes (Fig. 6B), indicating that Nf1^Myf5^ MPs are hypersensitive to Notch pathway activation upon external ligand stimulation.

Placement of Nf1^Myf5^ p14 FACS-isolated MPs cultured for 2 d in proliferation medium on uncoated dishes showed increased proliferation (Fig. 6C, D), as seen for primary myoblasts before (Supplementary Fig. 6B). We thus assessed, whether Jagged-1 treatment could reduce MP proliferation. Cultivation on Jagged-1-coated dishes (5 ng/μl) led to a reduced proliferation rate already in control p14 MPs, which was exacerbated in Nf1^Myf5^ MPs (Fig. 6C, D). This was confirmed by a dose-response titration using primary myoblasts showing that 2,5 ng/μl Jagged-1 reduced Nf1^Myf5^ myoblast proliferation rates to control levels, while higher concentrations reduced Nf1^Myf5^ myoblast proliferation rates below control levels (Supplementary Fig. 6E).

Both control and Nf1^Myf5^ MPs had lost Pax7 expression after 2 d cultivation on control dishes (Fig. 6C, E) consistent with previous observations (Liu et al., 2013). Jagged-1 maintained Pax7 expression in control and Nf1^Myf5^ MPs (Fig. 6C, E), with a relative increase in Pax7 abundance in Nf1^Myf5^ MPs (Fig. 6E). Conversely, Jagged-1 reduced the relative numbers of MyoD^+^ cells (Fig. 6F, G) and MyoD abundance (Fig. 6F, H) in control MPs, which was both exacerbated in Nf1^Myf5^ MPs (Fig. 6F-H), indicating that Jagged-1 induces a shift toward quiescence in juvenile control MPs, which is intensified in Nf1^Myf5^ MPs.

In Nf1-deficient oligodendrocytes, a Mek/Erk/nitric oxide synthase (NOS)/cyclic guanosine monophosphate (cGMP)/protein kinase G (PKG) pathway drives Notch pathway activation (Lopez-Juarez et al., 2017). GSEA showed NO-cGMP signaling enriched in Nf1^Myf5^ MPs (Fig. 6I). Consistent with this, Nf1^Myf5^ MPs placed on Jagged-1-coated dishes and treated with Mek inhibitor UO126 or pan-NOS inhibitor L-NAME cancelled the hyper-responsiveness to Jagged-1 (Fig. 6J).

We conclude that in juvenile MPs, a Ras/Mek/Erk/NOS pathway funnels into activation of the Notch pathway, inducing quiescence, which is exacerbated by lack on Nf1.

### Inhibition of Notch signaling prevents quiescence shift of Nf1^Myf5^ juvenile MPs and ameliorates the whole-body phenotype of Nf1^Myf5^ mice

Notch signaling regulates cell metabolism in several systems (Adams and Jafar-Nejad, 2019; Aster et al., 2017; Bi and Kuang, 2015). We thus first analyzed whether Notch signaling is upstream of metabolic gene expression in juvenile MPs. RT-qPCR of selected glycolysis and mitochondrial gene expression levels in WT p14 MPs cultured on control dishes or in the presence of Jagged-1 indicated that the Notch pathway can inhibit energy metabolism-related gene expression in juvenile MPs (Fig. 7A). Jagged-1 stimulation especially affected glycolytic genes as *Pfkfb1, Pfkfb3, Pfkm, Eno3, Ldha* and *Hk2*, and it mildly affected *mtCO1* and *Ndufv1* as representatives of the TCA cycle and ETC (Fig. 7A), overlapping transcriptome data of Nf1^Myf5^ p7 MPs. This suggests that in juvenile MPs, activation of the Notch pathway contributes to metabolic reprogramming by inhibiting glycolytic gene expression.

**Fig. 7.**
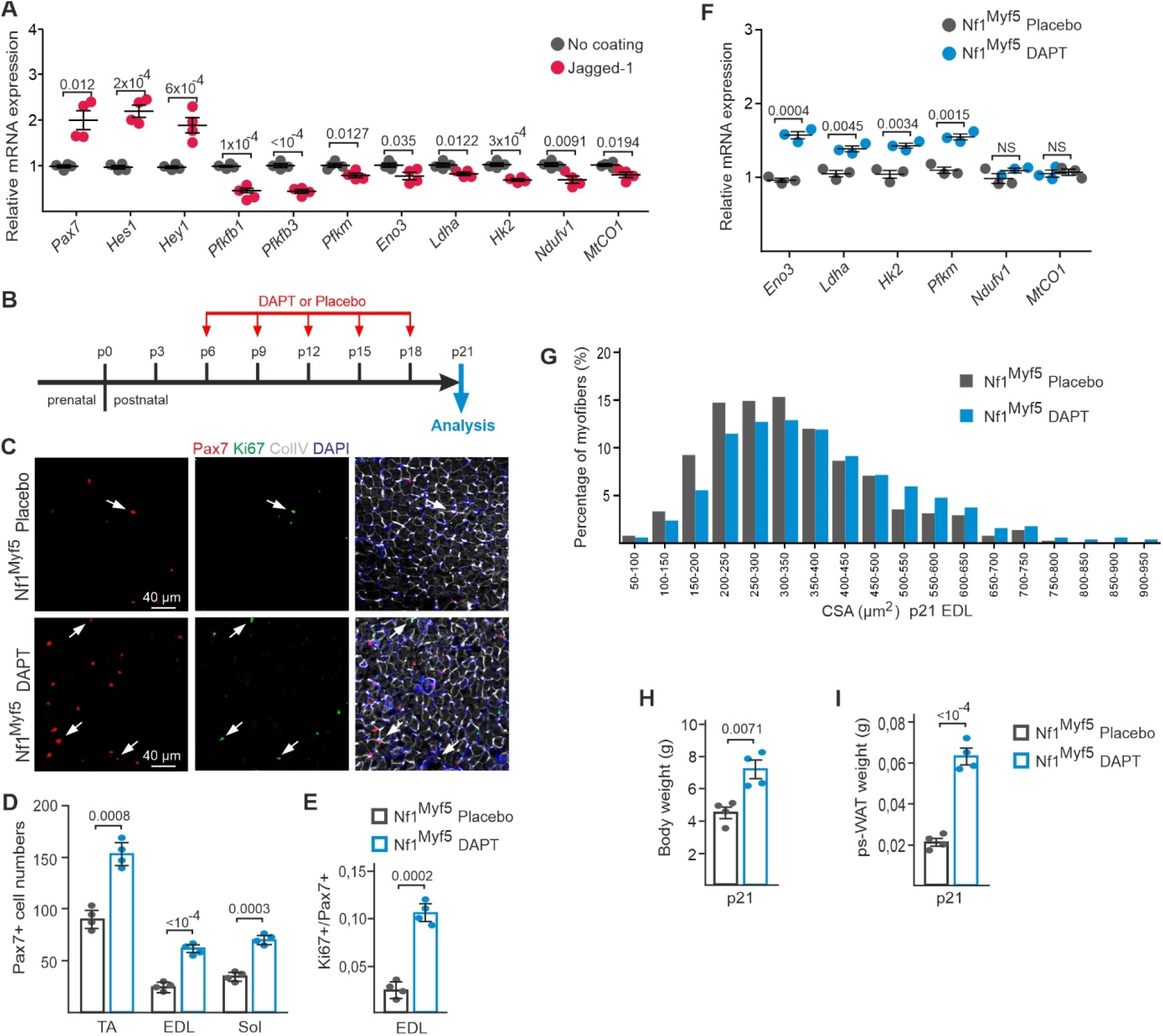
Rescue of Pax7 cell depletion, cell cycle exit, and metabolic reprogramming by Notch pathway inhibition. **A)** RT-qPCR of selected glycolysis, TCA, and OXPHOS genes on FACS-isolated WT p14 MPs cultured on Matrigel without coating or Jagged-1 coating for 48 h. *Pax7, Hes1*, and *Hey1* tested as internal controls (*n* = 4; each dot represents the mean of three replicates from one sample). **B)** Schematic depiction of DAPT treatment. **C)** Representative images of postnatal Nf1^Myf5^ mice sections treated with placebo or DAPT, stained for Pax7, Ki67, collagen IV, and DAPI. **D)** p21 Pax7^+^ cell quantification in Nf1^Myf5^ mice treated with placebo or DAPT (*n* = 4). **E)** p21 Ki67^+^/Pax7^+^ cell quantification relative to Pax7^+^ cells in Nf1^Myf5^ mice treated with placebo or DAPT (*n* = 4). **F)** RT-qPCR for glycolysis, TCA, and OXPHOS genes on muscle tissue from control or Nf1^Myf5^ mice treated with placebo or DAPT (*n* = 3; each dot represents the mean of three replicates from one sample). **G**) Distribution of myofiber diameter in control and Nf1^Myf5^ mice treated with placebo or DAPT (*n* = 4). **H)** Body weight of control or Nf1^Myf5^ mice treated with placebo or DAPT (*n* = 4). **I)** Posterior subcutaneous white adipose tissue weight in control and Nf1^Myf5^ mice treated with placebo or DAPT (*n* = 4). Data are mean± SEM; *P*-value calculated by two-sided unpaired *t*-test; *N*-numbers indicate biological replicates (mice per condition).

To test whether Notch signaling hyperactivation in Nf1^Myf5^ MPs is needed for premature quiescence induction and long-term metabolic reprogramming *in vivo*, we treated Nf1^Myf5^ pups with 5 doses of 30 mg/kg DAPT (or placebo control) from p6 to p18 (Fig. 7B).

To monitor Notch pathway inhibition *in vivo*, we measured the expression of two Notch pathway components and four Notch pathway targets using RT-qPCR and found reduced expression of all genes analyzed (Supplementary Fig. 7). This indicated that aberrant Notch pathway activity was not abrogated but effectively blunted by our DAPT regime in Nf1^Myf5^ mice. *In vivo* DAPT treatment increased Pax7^+^ cell numbers (Fig. 7C, D) and Pax7^+^ cell proliferation (Fig. 7C, E) compared to placebo-treated Nf1^Myf5^ mice. Thus, premature quiescence induction was prevented by *in vivo* DAPT treatment. *In vivo* DAPT treatment increased expression of glycolytic genes in p21 Nf1^Myf5^ muscle, while *Ndufv1* and *mtCO1* expression stayed the same (Fig. 7F).

Nf1^Myf5^ mice show muscle atrophy and a whole-body catabolic phenotype with attrition of white adipose tissue because of increased muscular consumption of fatty acids (Wei et al., 2020). *In vivo* DAPT treatment increased the apparent myofiber size (Fig. 7G) and the body weight (Fig. 7H) of Nf1^Myf5^ mice, indicating partial rescue of the muscle growth phenotype. In addition, *in vivo* DAPT treatment of Nf1^Myf5^ mice increased the white adipose tissue depot weight (Fig. 7I), indicating rescue of aberrant myofiber lipid metabolism.

We conclude that the Notch pathway is needed *in vivo* to induce premature quiescence in Nf1^Myf5^ MPs, and inhibition of the Notch pathway ameliorates metabolic reprogramming and improves the whole-body catabolic phenotype of Nf1^Myf5^ mice.

## Discussion

Early postnatal myofiber growth depends on a combination of cell accrual and metabolic growth. In addition to reduced metabolic growth (Wei et al., 2020), Nf1^Myf5^ muscle shows highly reduced myonuclear accrual during early postnatal growth.

### Nf1 counteracts Notch-dependent postnatal quiescence induction

To enable postnatal muscle growth by myonucelar accrual, a Pax7^+^ MP pool is maintained in proliferation, from which cells continuously differentiate and fuse to preexisting fibers. This, however, needs to be coordinated with quiescence induction in a subset of cells, which is required to establish a permanent MuSC pool. In Nf1^Myf5^ mice, postnatal MPs show premature quiescence induction evidenced by precocious and lasting cell cycle exit, reduced differentiation capability, reduced cell size and mTORC1 activity, and a transcriptomic signature showing increased expression of quiescence-related and reduced expression of differentiation-related genes. Therefore, the intricate balance between cell cycle exit/quiescence induction and maintenance of the progenitor pool and concomitant differentiation is distorted, leading to precocious contraction of the Pax7^+^ pool and so decreasing the numbers of both, myonuclei and MuSCs.

Quiescence induction is concomitant with changes in the epigenetic landscape. Reduced global H3K27me3 levels in Nf1^Myf5^ MPs overlap reduced H3K27me3 levels in quiescent adult MuSCs (Liu et al., 2013; Machado et al., 2017). Reduced H3K27me3 levels at the Pax7 locus and consistently increased mRNA and protein expression might support a shift to quiescence (Machado et al., 2017; Olguin and Olwin, 2004). Global H3K4me3 levels do not highly change in Nf1^Myf5^ MPs, also resembling adult MuSCs (Liu et al., 2013). However, early activation of MuSCs causes an increase in H3K4me3 levels at myogenic loci, including *Mylk2* (Machado et al., 2017), while in Nf1^Myf5^ MPs, this locus showed reduced H3K4me3 decoration (Supplementary Fig. 5F).

Nf1^Myf5^ juvenile MPs show an increase in hypermethylated DNA regions, including myogenic genes such as *Myl1*. The hypermethylation is consistent with previous studies showing increased DNA methylation in quiescent MuSCs (Carrio et al., 2015; Laker and Ryall, 2016) and increased heterochromatin content of quiescent MuSCs (Boonsanay et al., 2016; Robinson and Dilworth, 2018). This suggests that postnatal quiescence induction follows a change in the epigenetic landscape that contrasts adult MuSC quiescence exit.

Deficiency of tumor suppressors such as *Nf1* is usually associated with increased cell proliferation (Ratner and Miller, 2015). Opposite to their *in vivo* behavior, Nf1^Myf5^ MPs show enforced proliferation *in vitro*, which fully depends on Erk signaling. In accordance, Mek/Erk signaling is needed for *in vitro* myoblast proliferation (Jones et al., 2001), and Erk promotes MuSC proliferation in isolated myofibers (Griger et al., 2017). In contrast, Erk1/2 signaling has also been involved in quiescence induction in myoblasts *in vitro* (Reed et al., 2007); Mek/Erk signaling downstream of Ang-1/Tie2 induces quiescence-related gene expression in MPs *in vitro*, and *in vivo* Ang-1 overexpression increases and Tie2 blockade reduces the number of quiescent MuSCs (Abou-Khalil et al., 2009). This altogether indicates that Mitogen-activated protein kinase (Mapk/Erk) signaling in MPs/MuSCs is time- and context dependent and might strongly depend on the local microenvironment.

Notch pathway activation via recombinant Jagged-1 completely reverses *in vitro* hyperproliferation of Nf1^Myf5^ juvenile MPs, and Jagged-1 stimulation of both Nf1^Myf5^ and WT MPs increases Pax7 expression, reduces proliferation, and reduces MyoD expression, thus recapitulating the *in vivo* behavior. Inhibition of Notch signaling *in vivo* prevents precocious cell cycle exit and rescues muscle growth and MuSC numbers in Nf1^Myf5^ mice. Nf1^Myf5^ MPs are hypersensitive to Notch ligand stimulation, indicating a synergistic action of Ras/Mek/Erk and Notch signaling in early postnatal MPs, which is also observed in other contexts (Chen et al., 2015; Tremblay et al., 2013). As in oligodendrocytes (Lopez-Juarez et al., 2017), Nf1/Ras/Mek/Erk and Notch pathways are interconnected in juvenile MPs by a conserved NOS-dependent pathway. Notch signaling is essential for the maintenance of adult satellite cell quiescence (Bjornson et al., 2012; Fukada et al., 2011; Mourikis et al., 2012b), and is involved in postnatal MPs quiescence induction (Fukada et al., 2011; Kim et al., 2016; Mourikis et al., 2012a) as well as return of injury-activated MuSCs to quiescence, thereby promoting cell cycle exit and Pax7 expression (Wen et al., 2012). Our data further support the role of Notch signaling in driving postnatal MP quiescence at the expense of transient amplifying pool amplification and differentiation. These findings indicate an intrinsic function for Nf1 controlling Ras/Mek/Erk-based Notch pathway intensification in juvenile MPs, preventing quiescence induction, allowing pool expansion, and safeguarding postnatal muscle growth.

### Metabolic reprogramming in juvenile MPs quiescence induction

KEGG analysis of Nf1^Myf5^ MP transcriptome data indicates a global downregulation of anabolic processes, and GO analysis of hypermethylated regions shows that biosynthetic processes, RNA synthesis and transcription are affected. This result is consistent with a global decrease in biosynthesis and transcriptional activity in quiescent stem cells (van Velthoven and Rando, 2019) and suggests the induction of a transcriptionally controlled state of metabolic quiescence in Nf1^Myf5^ MPs. Adult MuSC activation is associated with an increase in glycolytic metabolism (Machado et al., 2017; Pala et al., 2018; Ryall et al., 2015; Ryall and Lynch, 2018), which is also seen in several other stem cell types, as glycolysis satisfies the proliferating cell’s demand for quick energy production and metabolite supply (Lunt and Vander Heiden, 2011; Ly et al., 2020). Most glycolytic genes are downregulated in Nf1^Myf5^ juvenile MPs, consistent with a highly reduced ECAR. Of these, *Pfkfb1* encodes one of four 6-phosphofructo-2-kinase/fructose-2,6-biphosphatase enzymes and is a key positive regulator of glycolytic flux (Atsumi et al., 2002). Together with the downregulation of genes encoding other key glycolysis pacemakers, such as *Hk2, Pfkm, Pkm*, and genes of the pyruvate dehydrogenase complex, this may explain the stalled glycolysis in Nf1^Myf5^ juvenile MPs. Notably, also genes of other carbohydrate metabolism pathways, such as the serine-one carbon and pentose phosphate pathways (*Phgdh, Psat1, Psph, G6pdx*; Supplementary Data 1) are downregulated, suggesting reduced folate metabolism and pentose production, both needed for nucleotide biosynthesis and cell division. Oxidative metabolism is, however, only mildly affected in p7 Nf1^Myf5^ MPs. Genes encoding ETC subunits or citrate cycle enzymes are mildly downregulated and hypermethylated, and genes for fatty acid metabolism are mostly unaffected. This is consistent with the idea that quiescent MuSCs mainly depend on low-level fatty acid–driven oxidative phosphorylation (OXPHOS) (Pala et al., 2018).

Notch signaling is highly context dependent and promotes glycolysis, for example, in hematopoietic stem cells or breast cancer cells (Ciofani and Zuniga-Pflucker, 2005; Landor et al., 2011) or represses glucose metabolism, for example, in the developing nervous system (Kuwabara et al., 2018) or mesenchymal stem cells (Lee and Long, 2018). Treatment of WT juvenile MPs with a Notch ligand strongly inhibits glycolytic gene expression and to a milder extent OXPHOS gene expression, indicating a direct role of Notch signaling in metabolic quiescence induction in MPs.

Reduced glycolytic flux increases cellular NAD^+^ levels, which activate Sirt1 in MuSCs, leading to deacetylation and inhibition of myogenic differentiation genes (Ryall et al., 2015). H4K16ac decoration is highly reduced in Nf1^Myf5^ juvenile MPs. Myogenic genes affected in Nf1^Myf5^ MPs overlap genes identified as Sirt1 targets in MuSCs (Ryall et al., 2015), indicating Sirt1-mediated metabolic reprogramming of Nf1^Myf5^ juvenile MPs, preventing myogenic differentiation. Therefore, inhibition of glycolytic genes downstream of Notch signaling, which is exacerbated by Ras/Mek/Erk input in Nf1^Myf5^ MPs, might be key for driving postnatal MP quiescence via metabolic reprogramming.

The main features of metabolic reprogramming are conserved in fully differentiated Nf1^Myf5^ muscle on a transcriptome level and functionally, with stalled glycolytic metabolism resulting in an energy deficit and muscle atrophy (Wei et al., 2020). Myofiber-specific *Nf1* inactivation has no deleterious consequences, indicating that metabolic perturbation of Nf1^Myf5^ muscle fibers is secondary to Nf1/Notch-mediated reprogramming of juvenile MPs, manifesting in long-term inhibition of glycolytic genes and persistently inhibiting carbohydrate metabolism. Thus, in a muscle-specific NF1 mouse model, the muscular metabolic phenotype can be fully traced back to *Nf1* activity in juvenile MPs. This model evidences transmission of differential gene expression signatures across cell divisions and cellular differentiation, which might involve epigenetic memory (Cavalli and Heard, 2019). Early-life Notch pathway inhibition ameliorates muscle defects and the whole-body catabolic state of Nf1^Myf5^ mice during the first three weeks of postnatal life. It will be interesting to see if this effect is persistent, or if treatment has to be repeated at later time points. Nevertheless, as the phenotype we observe in Nf1^Myf5^ animals critically develops within the first weeks of life we propose that the efficacy of treatment targeting Nf1-dependent pathways towards improvement of muscular function may depend on a critical early postnatal time window.

## Materials and Methods

### Animals and animal study approval

Mouse lines used in this study were described before; Myf5^Cre^ (Tallquist et al., 2000), Lbx1^Cre^ (Vasyutina et al., 2007), Nf1^flox^ (Zhu et al., 2001), Acta1^Cre^ (Miniou et al., 1999), and Rosa26^mTmG^ (Muzumdar et al., 2007). All animal procedures conducted within this study have been approved by the appropriate ethics committee and have therefore been performed in accordance with the ethical standards laid down in the 1964 Declaration of Helsinki and its later amendments. Animal studies were conducted in accordance with FELASA guidelines and were approved by the responsible authority (Landesamt für Gesundheit und Soziales Berlin, LaGeSo) under license numbers ZH120, G0346/13, G0176/19 and G0270/18. Timed matings were set up, and mice were sacrificed by cervical dislocation, fetuses were sacrificed by decapitation.

### Extraction of primary cells

Muscles from both fore- and hind limbs were minced and digested with 2 mg/ml collagenases A (Sigma-Aldrich, #11088793001) at 37 °C, for 1.5 hours. Tissue was further homogenized with a syringe; cells were washed cells twice with FACS buffer (PBS supplemented with 2% FBS, 1mM EDTA) and centrifuged with 300 g for 5 min at 4 °C. Tissue aggregates were removed using a 70 µm cell strainer (Miltenyi Biotec). Cells were resuspended in FACS buffer and incubated with antibodies: CD45 (Thermo Fisher, # 17-0451-83; 1:100), CD31 (Thermo Fisher, # 17-0451-83; 1:100), Sca-1 (BioLegend, # 108126; 1:200), Ter119 (Thermo Fisher, # 17-5921-83; 1:100) and α7-Integrin (R and D Systems, # MAB3518; 1:200) on ice for 30 min. This was followed by centrifugation with 300 g for 5 min at 4 °C, and washing with FACS buffer. Cell sorting was performed sing FACS Aria II SORP (BD Biosciences). Propidium Iodide (eBioscience, # 00-6990-50; 1:1000) was used to evaluate cell viability. Myogenic progenitors were gated for α7-Integrin after eliminating all CD45, CD31, Sca-1, and Ter119 positive cells from all mononuclear cells. FACS-sorted cells were harvested into tubes containing FACS buffer (for RNA or gDNA isolation, and for cytospin) or proliferation medium for culture (DMEM with 20% FBS, 1mM L-Glutamine, 1x Penicillin/streptomycin).

FACS-isolated cells were centrifuged with 500 g for 5 min, re-suspended in proliferation medium and counted using an automated cell counter (Luna™). Desired cell numbers were seeded on 12 mm coverslips that were counted with Matrigel (10 % Matrigel (Corning, # 356231) for 30 minutes). Cells were kept in proliferation medium, or immediately after adhesion subjected to differentiation conditions (DMEM with 2% HS, 1mM L-Glutamine, 1x Penicillin/streptomycin).

Primary myoblasts were isolated using pre-plating. After digestion (as above), the single cell solution was put on plastic dishes for 2 h to separate adherent fibroblast. Supernatant was taken off and transferred to dishes coated with 0.1% gelatin (Sigma-Aldrich, # G1393). Cells were cultured in proliferation medium for three days and detached using 0.25 % trypsin-EDTA (Thermo Fisher, #11560626). Cells were again transferred to uncoated plastic dishes, left for 45 min, and supernatant was transferred to gelatin coated dishes.

### Single muscle fiber isolation and processing

EDL muscles from 15 week old mice were isolated and digested in 2 ml of collagenase solution (0.2 % collagenase type I in DMEM) at 37 °C in a water bath. During digestion the muscle was regularly checked to avoid over-digestion. Digestion was stopped by carefully transferring the muscle to a pre-warmed Petri dish. A large bore glass pipette was used to release single myofibres under a microscope. To re-equilibrate the medium during the procedure, every 10 min muscles were transferred to 37 °C and 5 % CO_2_ for 5 min. Fibers were fixed in pre-warmed 4% PFA/PBS for 5 min followed by washing in PBS for 10 min for 3 times. Fibers were incubated in 1% glycine in PBS to minimize PFA background staining. Fibers were permeabilised with 0.5% Triton X-100 in PBS for 10 min. For blocking, TSA blocking solution (Roche) was used at 4 °C overnight. Fibers were washed once in PBS for 5min and incubated anti-Myosin, Skeletal, Fast (Sigma-Aldrich# M1570, 1:500) for 1 h at room temperature. Fibers were washed for 5 min for three times in PBS followed by incubation with secondary antibody and DAPI in staining solution (1 x PBS, 0.03 % horse serum, 0.003 % BSA, 0.001 % Triton X-100) for 1 h at room temperature. Fibers were washed in PBS for 5 min for three times. Then, fibers were individually transferred to a glass slide and mounted with Fluoromount G. Fibers were visualized and photographed using an Axiovert 200M (Zeiss) equipped with AxioVision 4.6 software (Zeiss).

### Immunolabeling of cells

For cytospin analysis, FACS-sorted MPs were kept at 4° C for 1 h followed by spinning at 50 g at 4° C for 5 min. 5 × 10^4^ cells were used per poly-L-lysine (1:100 dilution with bidest, Millipore, # A-003-E) coated 10 mm coverslip. MPs from cell culture were washed once with PBS to remove medium. Cells were fixed with 4% PFA (Merck, #104005) in PBS at room temperature for 10 min, permeabilized with PBX (0.3% Triton X-100) at room temperature for 10 min, and blocked with 5% BSA (Carl Roth, # 8076) in PBX at room temperature for 1h. Primary antibodies were diluted in 5% BSA and cells were incubated at 4°C overnight. After washing with PBX for 10 min three times, cells were incubated with secondary antibodies and DAPI (Invitrogen, #62248, 1:1000) diluted in PBX for 1 h at room temperature. Cells were washed 10 min for three times with PBX, and coverslips were mounted using Fluoromount-G (Southern Biotech, # 0100-01), and visualized using LSM700 confocal microscope (Zeiss) with ZEN imaging software (Zeiss). The primary antibodies and dilutions used were: Mouse anti-Pax7 (DSHB # Pax7; 1:25), Goat anti-Pax7 (provided by C. Birchmeier; 1:100), Rabbit anti-Ki67 (Abcam # ab16667; 1:500), Mouse anti-Ki67 (BD Biosciences #550609; 1:500), Rabbit anti-MyoD (Cell Signaling Technology #13812; 1:300), Mouse anti-MyoD (BD Biosciences # 554130; 1:500), Mouse anti-MF20 (DSHB # MF 20; 1:50), Anti-Myosin Skeletal, Fast (Sigma-Aldrich # M1570; 1:500), Rabbit anti-pErk1/2 (Cell Signaling Technology # 9101; 1:300), Rabbit anti-phosphor (s235/236)-S6 (Cell Signaling Technology # 4858; 1:300), Goat anti-Desmin (R&D Systems # AF3844; 1:500), Anti-trimethyl-Histone H3 (Lys27) (Millipore # 07-449; 1:500), anti-acetyl-Histone H4 (Lys16) (Millipore # 07-329; 1:500) (see also **Supplementary Table 4**). Relative fluorescence intensity (RFU) was measured with ImageJ software.

### Immunolabeling of tissue sections

Mouse hind limbs were dissected and embedded with tragacanth (Sigma-Aldrich #G1128) on a cork plate followed by freezing in isopentane / dry ice for 10 seconds. Samples were stored at -80 °C. 10 μm sections (Microm HM355S) were used for all applications. For Pax7 staining, slides were fixed with pre-cooled methanol at -20 °C for 10 min, followed by antigen retrieval with antigen retrieval solution (2 mM EDTA) for 10 min at 95 °C in a water bath (Julabo). For all other antibodies, slides were fixed with PFA at room temperature for 10min. Slides were blocked with blocking buffer (5% BSA in PBX) for 1 h at room temperature. Primary antibodies were diluted in blocking buffer and slides were incubated overnight at 4 °C. For Pax7 staining, this was followed by at room temperature for 4 h. Primary antibody solution was removed, and slides were washed for 10 min for four times in PBX. Slides were incubated with secondary antibodies diluted in PBX for 1 h at room temperature, followed by washing with PBX for 10min for four times. Slides were mounted with Fluoromount-G (Southern Biotech, # 0100-01) and visualized using LSM700 confocal microscope (Zeiss) with ZEN imaging software (Zeiss). The primary antibodies and dilutions used were: Goat anti-Collagen IV (Millipore # AB769; 1:500), Rabbit anti-Laminin (Sigma-Aldrich #L9393; 1:500), Mouse anti-Pax7 (DSHB # Pax7; 1:10), Rabbit anti-Ki67 (Abcam # ab16667; 1:400), anti-acetyl-Histone H4 (Lys16) (Millipore # 07-329; 1:400), Mouse anti-β-Tubulin III (Sigma-Aldrich # T8578; 1:100) (see also **Supplementary Table 4**). TUNEL staining on tissue sections was performed using the DeadEnd™ kit (Promega) according to the manufacturer’s instructions.

### Immunoblotting

For protein isolation, homogenization was performed with using TissueLyser (Qiagen) with RIPA buffer (50 mM Tris-Hcl, pH 8.0; 150 mM Nacl; 1% NP-40; 0.5% Sodium deoxycholate; 0.1% SDS). Protein concentration was measured using the Pierce BCA Protein Assay Kit (Thermo Fischer #23225). Total protein was separated with SDS-PAGE gels and transferred to PVDF membrane (GE Healthcare). Membrane were blocked with 5% BSA in TBST for 1 h at room temperature. Primary antibodies were diluted in blocking buffer, and membranes were incubated overnight at 4 °C room. After washing in PBST for 10 min for 3 times, HRP-conjugated secondary antibodies were applied for 1 hour in PBS at room temperature. Images were acquired using a Fusion FX spectra gel documentation system (Vilber) with FUSION FX software. Primary antibodies used were: Rabbit anti-phosphor (Thr 389)-p70s6k (Cell Signaling Technology # 9205; 1:1000), Rabbit anti-pErk1/2 (Cell Signaling Technology # 9101; 1:1000), Rabbit anti-Erk1/2 (Cell Signaling Technology # 9102; 1:1000), Mouse anti-ß-Actin (Cell Signaling Technology # 58169; 1:1000), Mouse anti-ß-Tubulin (Sigma-Aldrich # T8328; 1:1000), Rabbit anti-Nf1 (Bethyl # A300-140A; 1:1000). Blots images were analyzed and relative protein level was calculated using the gray value measurement tool in ImageJ.

### Seahorse XF metabolic flux analysis

Seahorse XF96 extracellular flux analyzer (Agilent) was used to measure the ECAR and OCR. The cartridge sensor was hydrated with 200 μl of calibration solution (Agilent) at 37 °C without CO_2_ overnight. FACS sorted MPs were plated on XF96 cell culture microplates coated with 10% Matrigel in warm assay medium (Agilent), the cell culture plate was centrifuged with 200 g for 5 min. and left 37 °C without CO_2_ for 45 minutes. Measurements were performed using the glycolysis stress test kit (Agilent) according to the manufacturer’s instructions. Seahorse Wave Desktop Software (Agilent) was used for data analysis.

### Jagged-1 ligand and inhibitor treatment

One day before the experiment, a 24 well plate was coated with 10% Matrigel at 37 °C for 30 min. Supernatant was removed and 100µl of Jagged-1 ligand (5ng/µl in PBS) was added (or only PBS for control plates), plates were left at room temperature overnight. Freshly FACS sorted p14 control and Nf1^Myf5^ muscle progenitors were cultured on Matrigel w/o coating or Jagged-1 coating for 48 hours; 3 × 10^5^ cells were used for each well. Cells were cultivated in proliferation medium without further treatment, or in the presence of Mek inhibitor UO126 (Promega # V1121; 10µM) or pan-NOS inhibitor L-NAME (Sigma-Aldrich # N5751; 2 mM). Total RNA was isolated from each sample followed by RT-qPCR analysis for Notch target genes and selected glycolysis, TCA and OXPHOS gene expression analysis.

### Notch Signaling Inhibitor Injection

5 doses of γ-secretase inhibitor DAPT solution (Sigma-Aldrich #D5942; 30 mg/kg in 95% corn oil/5% ethanol) were applied by subcutaneous injection to four Nf1^Myf5^ pups from postnatal day 6 to p21. Four Nf1^Myf5^ pups injected with Placebo were used as controls. Animals were sacrificed at p21, and hind limb tissue was used for RNA extraction / RT-qPCR analysis, or cryosectioning.

### RT-qPCR

Total RNA was isolated using the RNeasy Micro Kit (Qiagen; tissue samples) or RNeasy Mini Kit (Qiagen; cells). cDNA was synthesized using 1 μg of mRNA using the SuperScript™ III Reverse Transcriptase (Invitrogen™ # 18080044) kit, Oligo(dT)20 primer (Invitrogen™ #18418020) and RNaseOUT™ Recombinant Ribonuclease Inhibitor (ThermoFisher Scientific # 10777019) according to manufactures instructions. Quantitative RT-PCR analysis was performed using the ABI Prism HT 7900 real time PCR detection system (Applied Biosystems) equipped with SDS software version 2.4 (ThermoFisher Scientific) using GOTaq qPCR Master Mix (Promega) or SYBR Green qPCR Master Mix (Life Technologies). Actb was used to normalize the expression of each gene, double delta Ct (ΔΔCt) method was used to calculate the relative expression level. All primers used were purchased from Eurofins Scientific and listed in **Supplementary Table 1**. RT-qPCR was performed on at least 3 biological replicates (individual animals or independent cell culture assays), and were performed in triplicates for each sample.

### RNA sequencing analysis

Total RNA from 5×10^5^ FACS sorted p7 MPs was isolated using the RNeasy Micro Kit (Qiagen). RNA quantity and quality was tested on a Qubit^®^Fluorometer (Invitrogen) and a Bioanalyzer 2100 (Agilent). Sequencing libraries was prepared following Roche’s “KAPA stranded mRNA Seq” library preparation protocol. 11 cycles of PCR were used for libraries amplification followed by sequencing with an Illumina HiSeq 4000 system in PE75bp mode. 45-72 million fragments were acquired in each sample. Mapping was performed using STAR 2.4.2a software with mouse genome (mm9). Read counts were generated with R Studio function Summarize Overlaps and normalized to RPKM based on the number of uniquely mapped reads. Differential expression analysis was performed with DESeq2 using default settings. Genes with an absolute fold change ≥ 2 and adjusted p-value ≤ 0.01 were used as significantly differentially expressed. GSEA analysis was performed with the entire gene list using GSEA software 4.0.1 desktop (Broad Institute). Curated KEGG gene set and all Gene ontology set from Molecular Signature Database (MSigDB) were used for over representation analysis. Web based DAVID 6.8 was used for functional annotation of differentially expressed genes. Heatmaps were generated using R Studio 3.4.3.

### ChIP-Sequencing (ChIPmentation)

1×10^5^ FACS sorted p7 MPs from one animal were used for each sample, and two samples were used for each genotype. Cells were fixed (150µl of PBS/2%FBS/1% Formaldehyde) for 10 min at room temperature. Cells were sonicated using a Bioraptor (Diagenode) in sonication buffer (10 mM Tris-Hcl pH 8.0, 0.25% SDS, 2mM EDTA, 1x Roche complete Protease Inhibitor (Roche # 11697498001) with setting “high”,15 x (30 sec on / 30 sec off). After short spin-down this was repeated. Sheared chromatin was cleared by centrifugation for 10 min at full speed at 4 °C, supernatant was transferred to a new 0.2 ml PCR stripe and 2 µl of antibody (Anti-acetyl-Histone H4 (Lys16), Millipore # 07-329; Anti-trimethyl-Histone H3 (Lys4) Millipore # 07-473; Anti-trimethyl-Histone H3 (Lys27) Millipore # 07-449) was added and incubates on a rotator at 4 °C overnight. Magnentic beads (Invitrogen #10003D or 10001D) were preparation by washing and blocking at 4 °C overnight with 0.1% BSA/RIPA buffer (see above). Chromatin was incubated with magnetic beads at 4 °C for 2 h. CHIPmentation reaction (Schmidl et al., 2015; Schworer et al., 2016) was performed by incubating beads with tagmentation buffer (12.5 µl 2 x TD buffer, 11.5 µl nuclease free water, 1µl Tn5 enzyme (Illumina #FC-131-1024)) at 37°C for 10min. De-crosslinking was performed by incubating beads with ChIP elution buffer (1% SDS, 100 mM NaHCO_3_, 250 mM NaCl) with proteinase K (0.5 mg/mL) (Sigma Aldrich # 70663) at 55 °C for 1 h, followed by incubation overnight at 65 °C. Samples were eluted using the MinElute kit (Qiagen #28004), followed by 16-18 cycles PCR amplification. AMPureXP beads (Beckman Coulter #A63881) were sued for library product size selection. Size selected library concentration was measured with the Qbit^®^Fluorometer (Invitrogen) using 1-2 ng/µl in a total volume of 13µl. Library quality was assessed using the Bioanalyzer2100 (Agilent), showing a fragment peak around 280bp. Primers used for amplification and barcoding of ChIPmentation libraries were purchased from Eurofins Scientific and are listed in **Supplementary Table 2**.

### ChIP-Seq Data Analysis

Illumina 2 × 75 base paired end reads where aligned to the mouse reference genome mm9 using bwa v0.7.15 (Li and Durbin, 2009). For all 3 histone marks, peaks where called using MACS2 version 2.1.2 (Gaspar, 2018) and annotated with overlapping and proximal genes in R with the ChIPpeakAnno library (Zhu et al., 2010). GSEA was performed as described for RNA seq. For genome browser tracks, reads per million [rpm] have been averaged over replicates. For TSS profile plots, normalized ChIP seq profiles +/- 10 kb around the TSS have been aligned and averaged.

### MeDIP-Sequencing

Genomic DNA from 1×10^6^ FACS sorted p7 MPs from one animal was isolated with the All Prep DNA / RNA / Protein Mini Kit (Qiagen) and used as one sample, and two samples were used for each genotype. Sample quality was measured using the Nanodrop 2000. The low input MeDIP protocol (Taiwo et al., 2012) was used for library preparation. 80ng of genomic DNA were fragmented to around 170bp followed by End Pair and A tailing with the End Pair mix (NEB Next Ultra DNA library preparation kit). After ligation of truncated TruSeq adapters with the NEB Next Ultra Ligation Module, samples were purified with the Ampure XP beads. Samples were mixed with IP buffer (MagMeDIP –kit; DIAGENODE) and denatured for 3 minutes at 95°C. Denatured DNA was incubated with anti-5-meC-antibody and prewashed magnetic beads over night at 4°C on a rotator. After capturing, beads were washed, and methylated fragments were incubated for 5 minutes at 55°C, and for 15 minutes at 100°C in a proteinase K/elution buffer mix (1 µl of proteinase K in 100 µl DIB-buffer; Diagenode). Capture efficiency was determined by qPCR against spiked-in Lambda-DNA fragments in precapture and postcapture library samples. Libraries were amplified in a final PCR step using barcoded TruSeq primers (Index prime; 12 cycles) and size selected on a 2 % agarose gel (230-320bp, peak at 282-294bp). Quality was assessed on an Agilent Bioanalyzer and library concentration was determined by Qbit and qPCR. Primers used for MeDIP Library preparation are shown in **Supplementary Table 3**

### MeDIP-Seq Data Analysis

Illumina 2×75 base paired end reads where aligned to the mouse reference genome mm9 using bwa v0.7.15 [v.s.] and analyzed with QSEA Bioconductor package v.1.12.0 (Lienhard et al., 2017). CpG enrichment profiles were calibrated using blind calibration method. Differentially methylated regions (DMRs) where called at 250 base windows at an FDR of 10%, and annotated with promoter (TSS +/- 500 bases), exonic, and gene body regions as well as model based CpG islands (CGI) (Wu et al., 2010) For GO term based over representation analysis, all DMRs overlapping a CGI where assigned to the next gene.

### Data deposition

Sequencing raw data are available at the Gene Expression Omnibus (GEO) database under the Super Series accession number GSE159026.

### Quantification and statistical analysis

Analysis of NGS experiments is outlined above. All other data quantification was based on at least three independent biological replicates (individual animals or independent assays), N-numbers are noted in the figure legends. From each sample, at least three technical replicates were performed. Error bars represent the standard error of the mean (SEM). Two-tailed Student’s t test with 95% confidence interval was used to evaluate the significance of differences between two groups. Quantification and alignments of NGS analysis for RNA-seq, ChIP-mentation and MeDIP are described in more detail in the methods section above.

## Supporting information

Sipplementary Figures and Tables

## Acknowledgments

We thank the animal facility of the Max Planck Institute for Molecular Genetics, Berlin for expert support; Carmen Birchmeier for providing mouse lines; Daniel M. Ibrahim for help with ChIP-Seq procedure; and Roswitha Merle (FU:stat team) for help with statistical analysis.

