## Supplementary material for "Neurofibromin 1 controls metabolic balance and Notch-dependent quiescence of juvenile myogenic progenitors": Sipplementary Figures and Tables

### Supplementary Figures and Tables

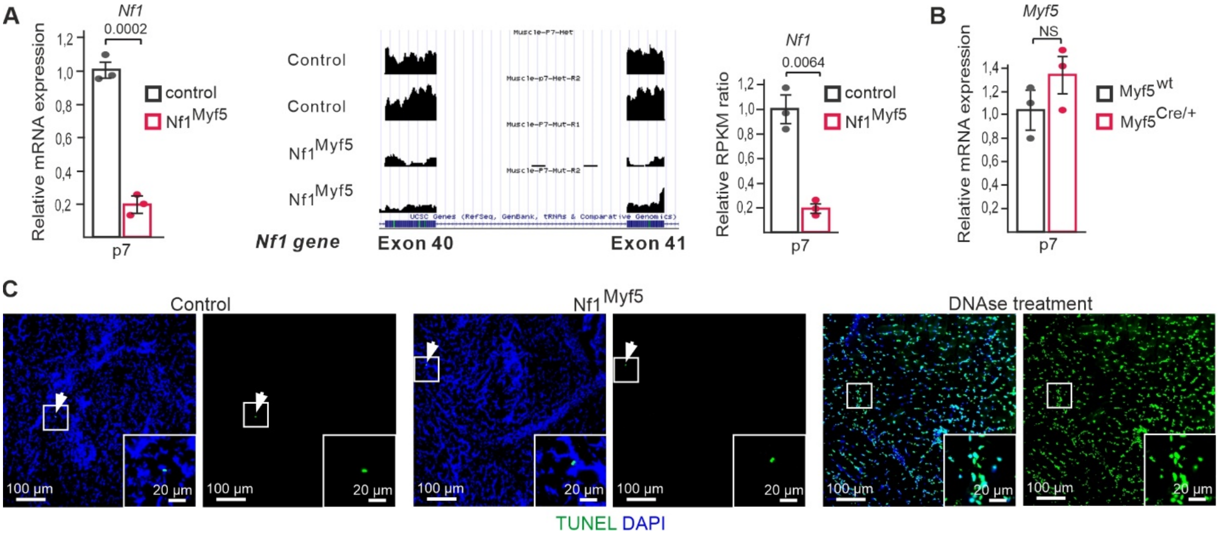


**Supplementary Fig. 1. Knockdown efficiency of *Nf1* and apoptosis analysis in Nf1$^{Myf5}$ mice**

**A)** Analysis of Nf1 knockdown efficiency in FACS isolated p7 MPs from control and Nf1$^{Myf5}$ mice by RT-qPCR (left; n=3 animals per genotype) and RNA-Sequencing (right, 2 animals per genotype). **B)** Analysis of *Myf5* expression in p7 MPs from Myf5$^{wt}$ vs. Myf5$^{Cre/+}$ animals (n=3 animals per genotype). **c)** Analysis of apoptosis assessed by TUNEL staining on p14 muscle sections. In both, controls as well as Nf1$^{Myf5}$ muscle, only occasional TUNEL+ cells can be seen. Right panel: positive control that was treated with DNAse.

Data are mean ± SEM; P-value calculated by two-sided unpaired t-test.

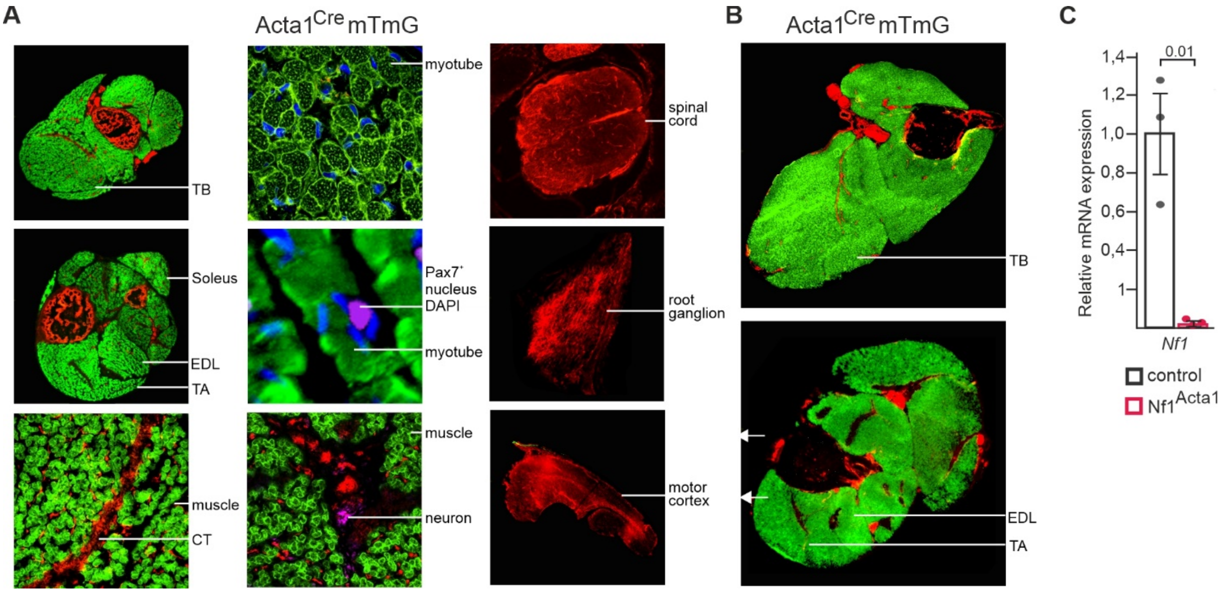


**Supplementary Fig. 2. Analysis of Acta1$^{Cre}$ efficacy**

**A, B)** Acta1-Cre mice were bred to Rosa26$^{mTmG}$ reporter mice and analyzed at embryonic day 18 (E18.5; a) or postnatal day 21 (p21; b). Staining: green depicts mG reporter activity, red depicts mT reporter activity. Satellite cells have been stained for Pax7, neurons for β-tubulin III (purple). **C)** RT-qPCR analysis of *Nf1* deletion efficacy in Nf1$^{Acta1}$ p21 muscle tissue (n=3 animals per genotype).

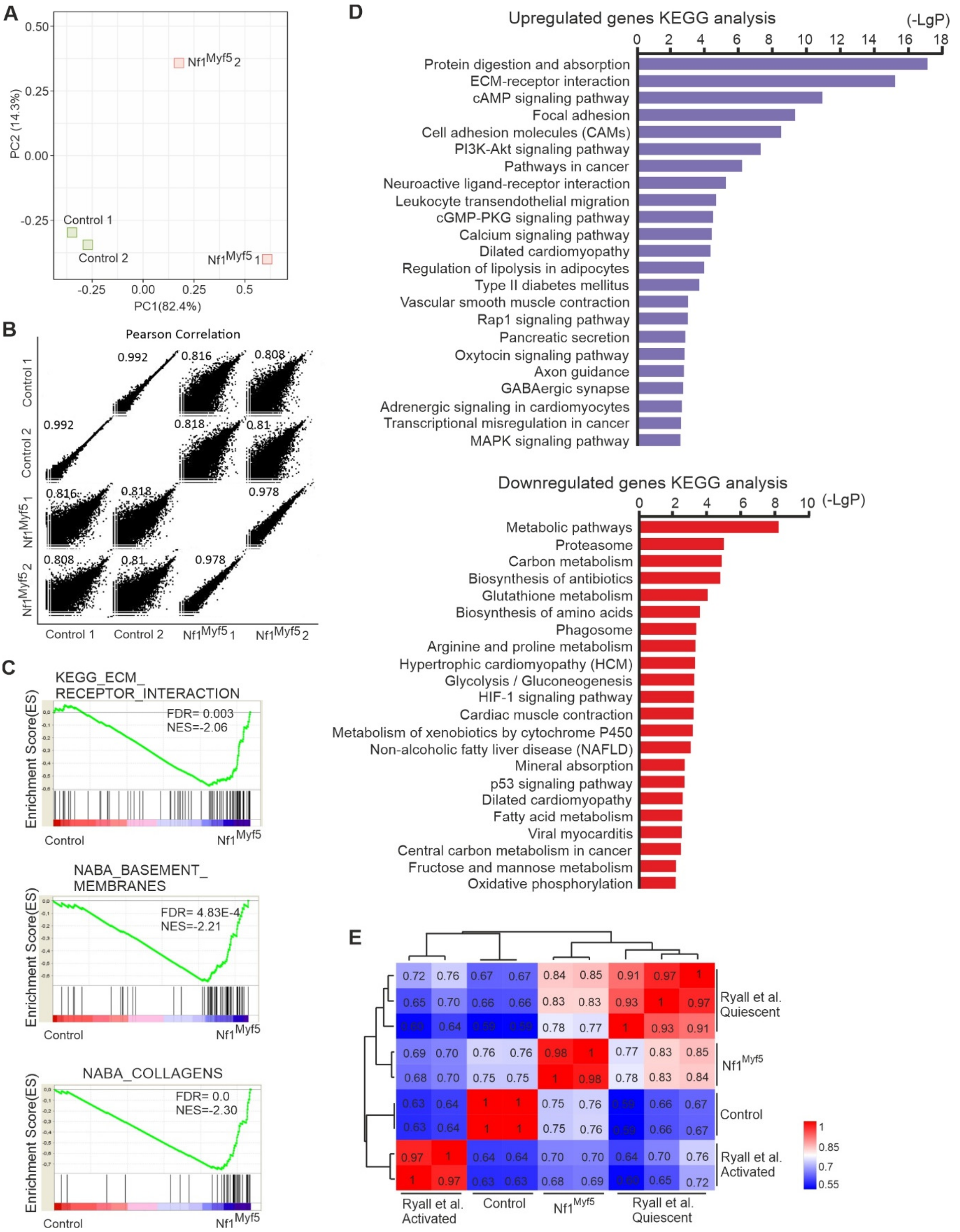


**Supplementary Fig. 3. RNA-Seq analysis of p7 Nf1$^{Myf5}$ MPs**

**A)** Principal component analysis of RNA-Seq data of control and Nf1$^{Myf5}$ p7 MPs. **B)** Pearson correlation analysis between RNA-Seq samples **C)** GSEA of control and Nf1$^{Myf5}$ p7 MP RNA-Seq data for “ECM Receptor Interaction”, “Basement Membranes” and “NABA Collagens”. **D)** KEGG analysis of genes downregulated (upper panel) or upregulated (lower panel) in Nf1$^{Myf5}$ p7 MPs. **E)** Correlation analysis of transcriptome data from Ryall et al. (2015) and transcriptomes of control or Nf1$^{Myf5}$ p7 MPs.

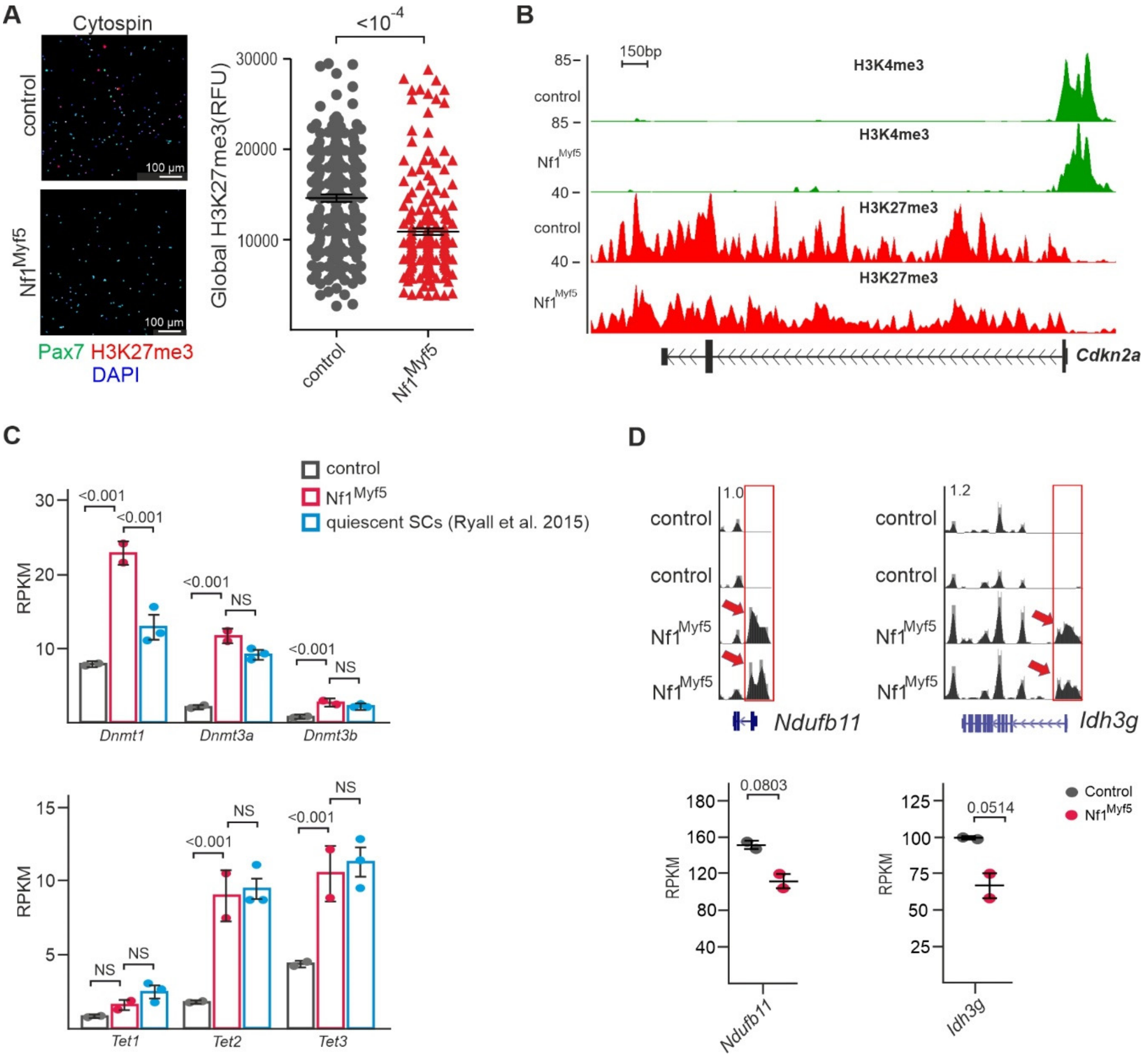


**Supplementary Fig. 4. H3K27me3 and DNA methylation analysis in Nf1$^{Myf5}$ MPs**

**A)** Immunolabeling for H3K27me3 on cytospun control and Nf1$^{Myf5}$ p7 MPs; quantification of fluorescence intensity is shown right (n=3 animals per genotype). **B)** ChIP-Seq tracks for H3K4me3 and H3K27me3 at the *Cdkn2a* locus in control and Nf1$^{Myf5}$ p7 MPs. **C)** Comparison of *Dnmt1*, *Dnmt3a*, *Dnmt3b*, and *Tet1-2* RPKM values between control / Nf1$^{Myf5}$ p7 MPs, and quiescent MuSCs from Ryall et al. (2015). **D)** Top row: MeDIP-Seq tracks from control and Nf1$^{Myf5}$ p7 MPs at the *Ndufb11* and *Idh3g* loci. RPKM values for *Ndufb1* and *Idh3g* expression levels from RNA-Seq of p7 MPs is shown below.

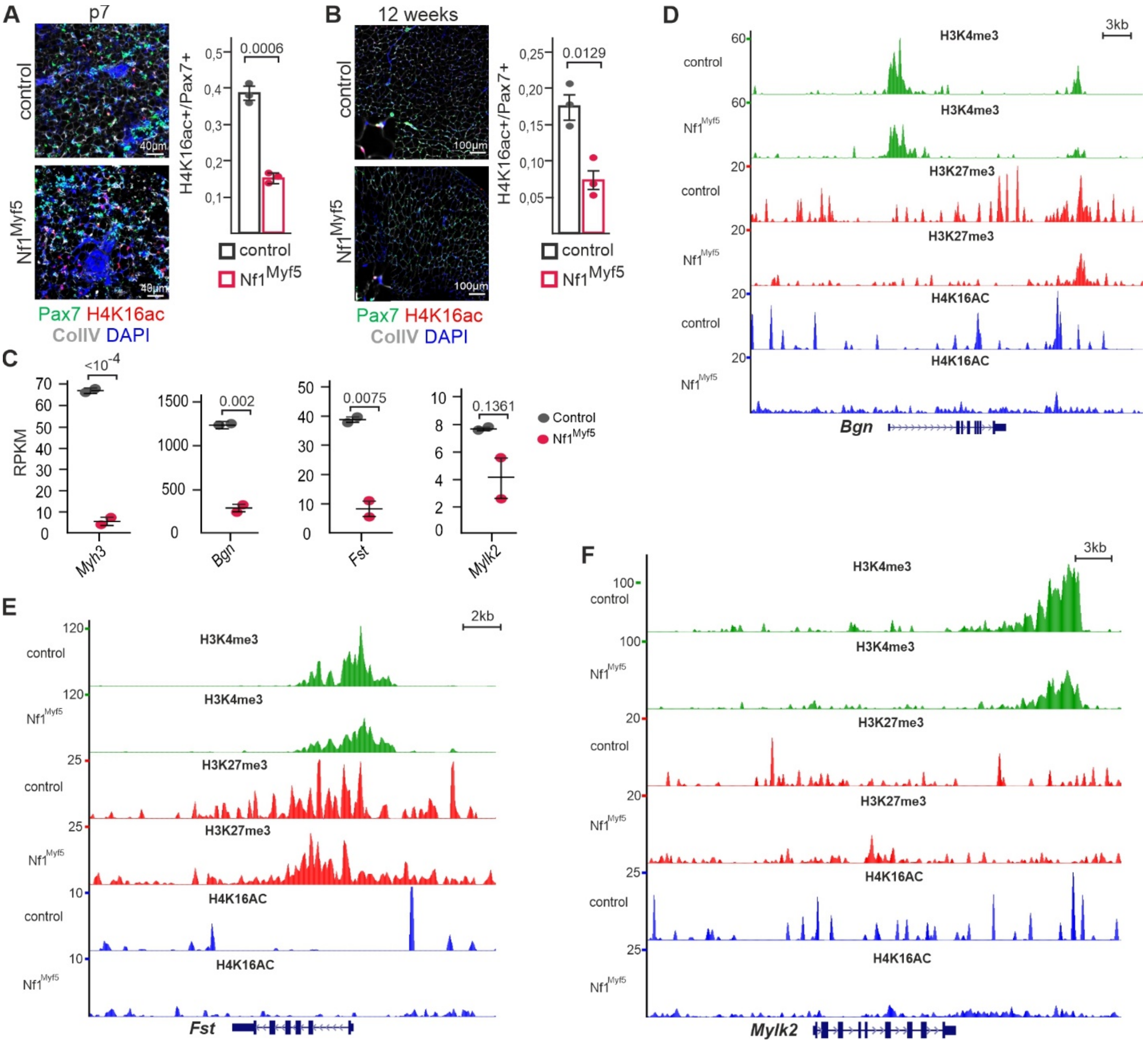


**Supplementary Fig. 5. Analysis of H4K16ac in Nf1$^{Myf5}$ mice**

**A, B)** Immunolabeling for Pax7 and H4K16ac on sections of TA muscles from p7 (b) and 12 weeks old (c) control and Nf1$^{Myf5}$ animals; quantification of H4K16ac+/Pax7+ cells relative to all Pax7+ cells is shown right (n=3 animals per genotype). **C)** RPKM values from RNA-Seq data of control and Nf1$^{Myf5}$ p7 MPs for *Bgn, Fst* and *Mylk2.* **D-F)** H3K4me3, H3K27me3, H4K16ac ChIP-Seq tracks from control and Nf1$^{Myf5}$ p7 MPs at the *Bgn, Mylk2 Fst* loci.

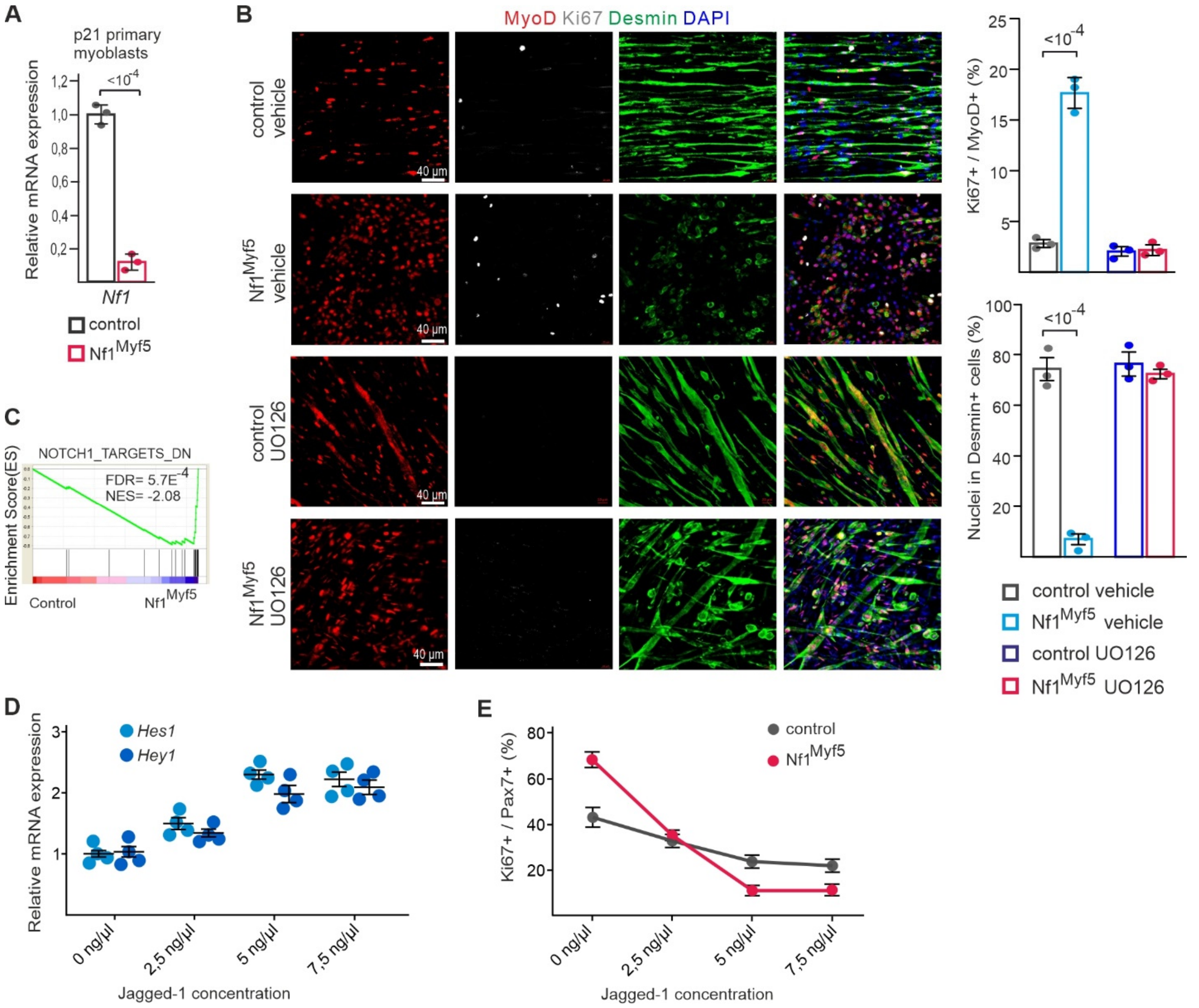


**Supplementary Fig. 6. Analysis of Nf1$^{Myf5}$ primary myoblasts**

**A)** RT-qPCR analysis of *Nf1* expression in p21 primary myoblasts from control and Nf1$^{Myf5}$ mice. **B)** Culture of control and Nf1$^{Myf5}$ p21 primary myoblasts immunolabeled for MyoD, Ki67 and Desmin. Cells were cultured in differentiation medium without addition, or with addition of MEK inhibitor UO126 for 2 days. Quantification of Ki67+ nuclei amongst MyoD+ nuclei, and quantification of nuclei in Desmin+ cells are shown below. **C)** GSEA on RNA-Seq data from control and Nf1$^{Myf5}$ p7 MPs for "Notch targets". Data are mean± SEM; P-value calculated by two-sided unpaired t-test. N-numbers indicate biological replicates (animals per genotype). **D)** Dose-response curve of indicated Jagged-1 ligand concentrations on Notch target gene (*Hes1*, *Hey1*) expression in wild type MPs. Expression levels in untreated MPs were set as 1. **E)** Dose-response curve of indicated Jagged-1 ligand concentrations on control and Nf1$^{Myf5}$ MP proliferation measured as numbers of Ki67+/Pax7+ cells.

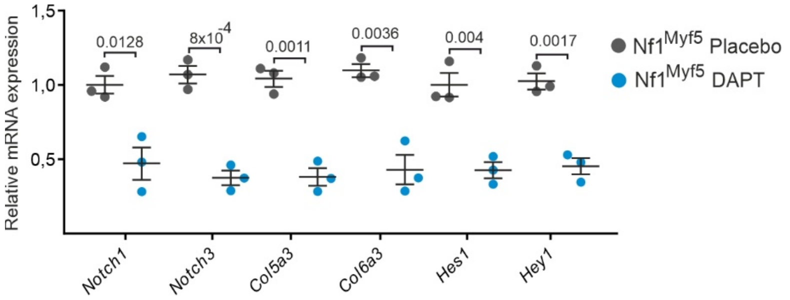


**Supplementary Fig. 7. DAPT treatment of $Nf1^{Myf5}$ animals**

RT-qPCR analysis of Notch pathway target gene expression in p21 TA muscle from control or $Nf1^{Myf5}$ mice treated with placebo or DAPT (n= 3 animals for each condition; each dot represents the mean value of three replicates from one sample). Data are mean ± SEM; *P*-value calculated by two-sided unpaired *t*-test).

#### Supplementary Table 1. Primer Sequences (RT-qPCR)

| | |
|---|---|
| Nf1_fw | ACAAAGGGTTACTGCCAT |
| Nf1_rev | GCTCCCCCAGATTTTTGC |
| Myf5_fw | TGAGGGAACAGGTGGAGAAC |
| Myf5_rev | CTGTTCTTTCGGGACCAGAC |
| Pax7_fw | CCGTGTTTCTCATGGTTGTG |
| Pax7_rev | GAGCACTCGGCTAATCGAAC |
| Spry1_fw | TAGGTCAGATCGGGTCATCC |
| Spry1_rev | TTCGCAGATGAACTTGTGCT |
| Notch1_fw | AGGCAAATGCCTCAACACAC |
| Notch1_rev | CATTGGAACTCCCCAATCTG |
| Notch3_fw | AGGGCCAGAACTGTGAAGTC |
| Notch3_rev | AGGGCACTGGCAGTTGTAAG |
| Hes1_fw | TCATCAAAGCCTATCATGGAGA |
| Hes1_rev | AGGTGCTTCACAGTCATTTCC |
| Hey1_fw | CCGACGAGACCGAATCAATA |
| Hey1_rev | TTTTCAGGTGATCCACAGTCA |
| Myod1_fw | AGCACTACAGTGGCGACTCA |
| Myod1_rev | GCTCCACTATGCTGGACAGG |
| MyoG_fw | CTACAGGCCTTGCTCAGCTC |
| MyoG_rev | AGATTGTGGGCGTCTGTAGG |
| Myh3_fw | GGACGCTGGAGGATCAAAT |
| Myh3_rev | AAAATGGATGCGGATGAACT |
| ATP2a1_fw | CTGACCGAAAGTCAGTGCAA |
| ATP2a1_rev | GGTGGATTTGATGGAGAGGA |
| DNMT1_fw | GAACCCCAGATGTTGACCAG |
| DNMT1_rev | GGTGTCGACAGGACACAGGT |
| DNMT3a_fw | ACTTGGAGAAGCGGAGTGAA |
| DNMT3a_rev | TTCTGGTGGGGTCTCAGTTC |
| Myl1_fw | ACAACAAGGACCAGGGAGGT |
| Myl1_rev | CTGCCAGCAACGCTTCTAC |
| Pfkfb1_fw | TGATGCCACCAACACTACCA |
| Pfkfb1_rev | CTGCAATGATGTCTGGGTCA |
| Calcr_fw | AGCCCAACTCCAGTTCTTCA |
| Calcr_rev | TCCTTCATAAGAGGGCAACTG |
| Pfkfb3_fw | GCAGTACAGCTCTTACAACTTCT |
| Pfkfb3_rev | TGTCCACCTTCCTTTGTCAG |

| | |
|---|---|
| Pfkm_fw | ACGTGACCAAGGCTATGGAT |
| Pfkm_rev | GACTGGGGGTCTGACATGAG |
| Eno3_fw | AAATCTTCGCCCGGGAAATC |
| Eno3_rev | TGCTTCATAGATACCCGTGGA |
| Ldha_fw | ACTTGGCGGATGAGCTTG |
| Ldha_rev | GCGGTGATAATGACCAGCTT |
| Hk2_fw | GTTTGACCACATTGCCGAAT |
| Hk2_rev | CACGCCACTGGACTTGAAC |
| Ndufv1_fw | CCA TTC TAA TCG CCATAG CC |
| Ndufv1_rev | GCC GTA TGG ACC AAC AAT G |
| Mtco1_fw | TGCTAGCCGCAGGCATTAC |
| Mtco1_rev | GGGTGCCCAAAGAATCAGAAC |
| Myh1_fw | CGGGAAGACTGTGAACACGA |
| Myh1_rev | CGTTCCCAAAGGCCTCCA |
| Myh2_fw | ACCCTCCCAAGTACGACAAG |
| Myh2_rev | TACACCGGCAGCCATTTGTA |
| Myh4_fw | CAGAGTCACCTTCCAGCTCA |
| Myh4_rev | TGATTTCACCTTGACTGACGT |
| Myh7_fw | AGCTGGGAAGACTGTCAACA |
| Myh7_rev | CCAAAGGCCTCCAGAGCA |
| Col5a3_fw | CCTGGGGTGTGCATAGAGAC |
| Col5a3_rev | GGACACTGAGAAGGCTGGAC |
| Col6a3_fw | GTGGCTCAGTATGCAGACACT |
| Col6a3_rev | GTCCAGAGAAGATCCCGTGT |
| Gapdh-fw | AACTTTGGCATTGTGGAAGG |
| Gapdh-rev | CAGTCTTCTGGGTGGCAGTG |
| Actb-fw | CGTGAAAAGATGACCCAGATCA |
| Actb-rev | GGGACAGCACAGCCTGGAT |

**Supplementary Table 2. Primer sequences (ChIP-Sequencing; Nextera custom primers)**

| | |
|---|---|
| Ad1noMX | AATGATACGGCGACCACCGAGATCTACACTCGTCGGCAG |
| Ad2.1_TAAGGCGA | CAAGCAGAAGACGGCATACGAGATTCGCCTTAGTCTCGT |
| Ad2.2_CGTACTAG | CAAGCAGAAGACGGCATACGAGATCTAGTACGGTCTCGT |
| Ad2.3_AGGCAGAA | CAAGCAGAAGACGGCATACGAGATTTCTGCCTGTCTCGT |
| Ad2.4TCCTGAGC | CAAGCAGAAGACGGCATACGAGATGCTCAGGAGTCTCGT |
| Ad2.5GGACTCCT | CAAGCAGAAGACGGCATACGAGATAGGAGTCCGTCTCGT |
| Ad2.6_TAGGCATG | CAAGCAGAAGACGGCATACGAGATCATGCCTAGTCTCGT |
| Ad2.7_CTCTCTAC | CAAGCAGAAGACGGCATACGAGATGTAGAGAGGTCTCGT |
| Ad2.8CAGAGAGG | CAAGCAGAAGACGGCATACGAGATCCTCTCTGGTCTCGT |
| Ad2.9_GCTACGCT | CAAGCAGAAGACGGCATACGAGATAGCGTAGCGTCTCGT |
| Ad2.10_CGAGGCTG | CAAGCAGAAGACGGCATACGAGATCAGCCTCGGTCTCGT |
| Ad2.11_AAGAGGCA | CAAGCAGAAGACGGCATACGAGATTGCCTCTTGTCTCGT |
| Ad2.12_GTAGAGGA | CAAGCAGAAGACGGCATACGAGATTCCTCTACGTCTCGT |

**Supplementary Table 3. Primer sequences (MeDIP Library preparation)**

| | |
|---|---|
| TRUSEQ_UNI_TRUNC | ACACTCTTTCCCTACACGACGCTCTTCCGATC*T |
| Index_SCAFFOLD | 5P-GATCGGAAGAGCACACGTCTGAACTCCAGTCAC |
| Index1 | CAAGCAGAAGACGGCATACGAGATCGTGATGTGACTGGAGTTCAGACGTGTGCTCTTCCGATC |
| Index8 | CAAGCAGAAGACGGCATACGAGATTCAAGTGTGACTGGAGTTCAGACGTGTGCTCTTCCGATC |
| Index10 | CAAGCAGAAGACGGCATACGAGATAAGCTAGTGACTGGAGTTCAGACGTGTGCTCTTCCGATC |
| Index11 | CAAGCAGAAGACGGCATACGAGATGTAGCCGTGACTGGAGTTCAGACGTGTGCTCTTCCGATC |
| 2$^{nd}$ primer | AATGATACGGCGACCACCGAGATCTACACTCTTTCCCTACACGACGCTCTTCCGATCT |
| 1CpG_qPCR_F | ACAAGTTGTTTGATCTTTGC |
| 1CpG_qPCR_R | CCTATGAGCAACGTGTTAG |
| 5CpG_qPCR_F | CACTTGAATCTGTGGTTCAT |
| 5CpG_qPCR_R | TAGAAAAAGACAACTCTGGC |
| 10CpG_qPCR_F | GAACTCACACACAACACCA |
| 10CpG_qPCR_R | ACTCTGAATACCGACTCAAT |
| 15CpG_qPCR_F | TATCACTGTTGATTCTCGC |
| 15CpG_qPCR_R | GGTAAAGAGTTTGGATTAGG |

#### Supplementary Table 4. Antibodies

| Antibody | Source | Identifier |
|---|---|---|
| Goat anti-Collagen IV | Millipore | Cat# AB769; RRID:AB_92262 |
| Goat anti-Desmin | R&D Systems | Cat# AF3844; RRID:AB_2092419 |
| Rabbit anti-Nf1 | Bethyl | Cat# A300-140A, RRID:AB_2149790 |
| Mouse anti-Pax7 | DSHB | Cat# pax7, RRID:AB_528428 |
| Guinea pig anti-Pax7 | This paper | N/A |
| Rabbit anti-Ki67 | Abcam | Cat# ab16667, RRID:AB_302459 |
| Mouse anti-Ki67 | BD Biosciences | Cat# 550609, RRID:AB_393778 |
| Rabbit anti-MyoD | Cell Signaling Technology | Cat# 13812, RRID:AB_2798320 |
| Mouse anti-MyoD | BD Biosciences | Cat# 554130, RRID:AB_395255 |
| Mouse anti-MF20 | DSHB | Cat# MF 20, RRID:AB_2147781 |
| Anti-Myosin (Skeletal, Fast) antibody | Sigma-Aldrich | Cat# M1570, RRID:AB_ 2147168 |
| Rabbit anti-phospho (Thr 389)-p70s6k | Cell Signaling Technology | Cat# 9205, RRID:AB_330944 |
| Rabbit anti- pERK1/2 | Cell Signaling Technology | Cat# 9101, RRID:AB_331646 |
| Rabbit anti- ERK1/2 | Cell Signaling Technology | Cat# 9102, RRID:AB_330744 |
| Phospho-S6 Ribosomal Protein (Ser235/236) | Cell Signaling Technology | Cat# 4858, RRID:AB_916156 |
| Anti-acetyl-Histone H4 (Lys16) | Millipore | Cat# 07-329, RRID:AB_310525 |
| Anti-trimethyl-Histone H3 (Lys4) | Millipore | Cat# 07-473, RRID:AB_1977252 |
| Anti-trimethyl-Histone H3 (Lys27) | Millipore | Cat# 07-449, RRID:AB_310624 |
| Mouse anti-β-Tubulin III | Sigma-Aldrich | Cat# T8578, RRID:AB_1841228 |
| Mouse anti-ß-Actin | Cell Signaling Technology | Cat# 58169, RRID:AB_2750839 |
| Mouse anti-ß-Tubulin | Sigma-Aldrich | Cat# T8328, RRID:AB_1844090 |
| Alexa Fluor 488 Donkey anti-Mouse | Thermo Fisher | Cat# A-21202, RRID:AB_141607 |
| Alexa Fluor 568 Donkey anti-Mouse | Thermo Fisher | Cat# A10037, RRID:AB_2534013 |
| Alexa Fluor 488 Donkey anti-Rabbit | Thermo Fisher | Cat# A-21206, RRID:AB_141708 |
| Alexa Fluor 568 Donkey anti-Rabbit | Thermo Fisher | Cat# A10042, RRID:AB_2534017 |
| Alexa Fluor 488 Donkey anti-Goat | Thermo Fisher | Cat#A-11055, RRID:AB_2534102 |
| Alexa Fluor 568 Donkey anti-Goat | Thermo Fisher | Cat# A-11057, RRID:AB_2534104 |
| Alexa Fluor 680 Donkey anti-Goat | Thermo Fisher | Cat# A-21084, RRID:AB_2535741 |
| Alexa Fluor 680 Goat anti-Guinea pig | Thermo Fisher | Cat# SA5-10098, RRID:AB_2556678 |
| HRP Goat anti-Rabbit | Thermo Fisher | Cat# A27036, RRID:AB_2536099 |
| HRP Goat anti-Mouse | Thermo Fisher | Cat# G-21040, RRID:AB_2536527 |
| CD31(PECAM1) APC | Thermo Fisher | Cat# 17-0311-82, RRID:AB_657735 |
| CD45 | Thermo Fisher | Cat# 17-0451-83, RRID:AB_469393 |
| Ter119 | Thermo Fisher | Cat# 17-5921-83, RRID:AB_469474 |
| Sca1 | BioLegend | Cat# 108126, RRID:AB_10645327 |
| Integrin alpha 7 | R and D Systems | Cat# MAB3518, RRID:AB_2128441 |
